# Lipid-mediated Association of the Slg1 Transmembrane Domains in Yeast Plasma Membranes

**DOI:** 10.1101/2021.06.29.450341

**Authors:** Azadeh Alavizargar, Annegret Elting, Roland Wedlich-Söldner, Andreas Heuer

**Affiliations:** Institute of Physical Chemistry, University of Muenster, Corrensstr. 28/30, 48149 Muenster, Germany; Institute of Cell Dynamics and Imaging, University of Muenster, Von-Esmarch-Str. 56, 48149 Muenster, Germany

## Abstract

Clustering of transmembrane proteins underlies a multitude of fundamental biological processes at the plasma membrane (PM) such as receptor activation, lateral domain formation and mechanotransduction. The self-association of the respective transmembrane domains (TMD) has also been suggested to be responsible for the micron-scaled patterns seen for integral membrane proteins in the budding yeast plasma membrane. However, the underlying interplay between local lipid composition and TMD identity is still not mechanistically understood. In this work we combined coarse-grained molecular dynamics (MD) simulations of simplified bilayer systems with high resolution live-cell microscopy to analyze the distribution of a representative helical yeast TMD from the PM sensor Slg1 within different lipid environments. In our simulations we specifically evaluated the effects of acyl chain saturation and anionic lipids head groups on the association of two TMDs. We found that weak lipid-protein interactions significantly affect the configuration of TMD dimers and the free energy of association. Increased amounts of unsaturated phospholipids strongly reduced helix-helix interaction, while the presence of anionic phosphatidylserine (PS) hardly affected dimer formation. We could experimentally confirm this surprising lack of effect of PS using the network factor, a mesoscopic measure of PM pattern formation in yeast cells. Simulations also showed that formation of TMD dimers in turn increased the order parameter of the surrounding lipids and induced long-range perturbations in lipid organization. In summary, our results shed new light on the mechanisms for lipid-mediated dimerization of TMDs in complex lipid mixtures.

## Introduction

The plasma membrane (PM) has to fulfill a wide range of biological functions such as selective uptake of nutrients and ions, signal transduction and cellular morphogenesis. For these diverse processes to be regulated efficiently, many resident proteins and lipids of the PM are laterally segregated into distinct functional domains. A striking example of such domains has been described for the budding yeast PM, where all integral membrane proteins as well as lipids form a patchwork of overlapping domains. ^1^ A systematic microscopic evaluation of yeast PM resident proteins demonstrated that domain formation depends on both the sequence of TMDs and the cellular lipid composition.^2^ Based on these findings, it was suggested that collective interactions between TMDs and lipids could drive the formation of a characteristic patchwork membrane. ^3^ A particularly useful observable to characterize PM patterns in yeast is the network factor (“intensity distribution” in ref. ^2^), which represents the area above cumulative intensity histograms for deconvolved total internal reflection microscopy (TIRFM) images. This parameter represents an intuitive measure for the heterogeneity in the lateral distribution of a specific PM marker. Diffuse arrangements that resemble network-like patterns yield large network factors, whereas strong clustering in patch-like structures results in low values. Thus, on a qualitative level the network factor can be regarded as a mesoscopic observable to characterize the effective association tendency of TMDs. However, a mechanistic understanding of lateral TMD association and segregation in the yeast PM is still lacking.

One way to approach this problem is to perform MD simulations of simplified mixtures of TMDs and lipids. Most TMDs consist of single *α*-helices with mostly hydrophobic or small amino acids. Many experimental techniques have been used to probe the factors involved in the association process of TMDs. These factors include the specific TMD sequence, the local lipid composition, binding of ligands to functional groups located at the extracellular or intracellular parts of the the TMD as well as general physico-chemical properties of the membrane. ^4^ Many TMDs contain specific sequence motifs. A well-studied example is the GxxxG dimerization motif, where x stands for hydrophobic residues with short or slightly polar side chains, ^5, 6^ that allow a steric match between interacting surfaces of TMDs. A more general ‘GxxxG-like’ motif has been proposed as SmxxxSm, where ‘Sm’ is a small residue (Gly, Ala, Ser or Thr).^7^ GxxxG-like motifs can influence both, the free energy of helix-helix association and the structural conformation of TMD dimers, characterized by specific crossing angles. ^4^ The biophysical properties of the lipid bilayer such as lipid order or hydrophobic thickness can have additional, non-specific effects on the association of TMDs. For example, highly ordered lipids in the gel phase exclude TMDs and thereby indirectly enhance local TMD concentration and dimerization. ^8^ On the other hand, increasing lipid order and the transition from Ld to Lo phase by the addition of cholesterol directly supports stronger association of TMDs.^9, 10^ Helix-helix association of TMDs is also affected by the hydrophobic thickness of the bilayer, ^11, 12^ which has also been studied computationally,^13–16^ or by combining simulations and experiments.^17^ In model membranes, increased bilayer thickness through addition of long acyl chains or cholesterol promotes the self-association of TMDs.^8^ Finally, dimerization of TMDs has been shown to be affected by either, the length and saturation of acyl chains in membrane lipids ^18–20^ or the charge of lipid headgroups. ^21^ MD simulations have also inspected the comparative dissociation free energy of different lipid types from TMD dimers. ^22, 23^ In addition to its non-specific effects through lipid order and membrane thickness, cholesterol has also been shown in coarse grained simulations to specifically interact with certain TMD stretches, including GxxxG or CRAC motifs, ^24^.^25^ Specific examples for this include the TMDs of ErbB2^26, 27^ or Glycophorin A (GpA).^19^ While bilayer propertiess and composition clearly regulate the association of TMDs, the association of proteins can in turn have large effects on the lipid environment. MD simulations have shown that a single TMD can already induce long-range perturbations in the hydrophobic core of the bilayer, mediating communication between TMDs at a distance of 2-3 nm from each other.^28^ Importantly, a reduction of perturbations on membrane thickness and membrane order has been observed upon helix-helix association, ^29^ leading to local variations in membrane thickness around dimers in model membranes with different lipid compositions. ^20^

Despite the large number of studies on selected TMD-membrane systems, the impact of weak, collective lipid interactions on helix-helix association as well as the complex interplay between dimer formation and local lipid organization is still insufficiently understood.

In this work, we combine computational and experimental approaches to study TMD-association in a simplified lipid bilayer system that recapitulates some features of the yeast PM. In order to select a representative TMD for our studies, we tested localization of isolated TMDs from various single-spanning integral yeast PM proteins fused to GFP. Of the TMDs that were correctly delivered to the PM, four contained the previously characterized GxxxG motifs — the cell wall stress sensors Mid2, Mtl1 and Slg1 and the cell fusion protein Fus1. We decided to focus on the Slg1 TMD, as it contained a total of three GxxxG motifs and one GxxxG-like motif (Figure 1A). The first two GxxxG motifs, called glycine zipper, have been previously recognized as a driving motif for homo- and oligomerization of membrane proteins. ^30–32^ In addition, the central GVxxGV motif has been identified as a strong interaction motif in the GpA TMD dimer ^7^ (Figure 1A). Finally, the cytosolic C-terminus of the Slg1 TMD contains two positively charged residues (Figure 1A) that might affect TMD association via electrostatic interaction with anionic lipid head groups.

**Figure 1:**
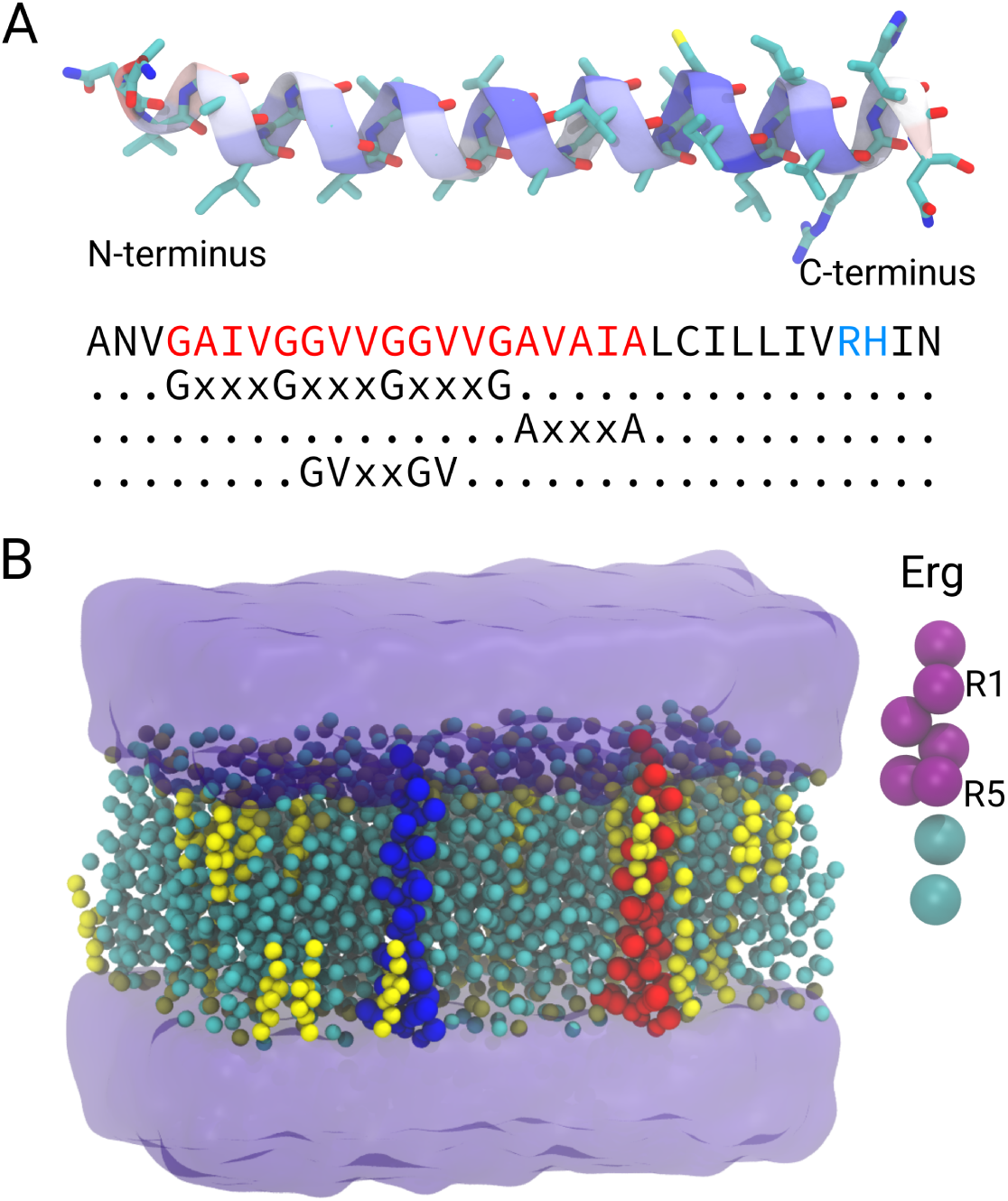
(A) The atomistic representation of the Slg1 TMD studied in this work, along with the associated sequence. The GxxxG and GxxxG-like motifs as well as the positively charged residues in the C-terminus are highlighted, respectively, in red and blue. (B) The simulations system is represented with the two TMD located at a distance of 4.5 Å from each other. The beads of the two TMDs and the lipids are shown in spheres (ergosterol molecules in yellow) and the water is shown in surface representation. Half of the simulation system is not shown for clarity. Ergosterol molecule is shown in MARTINI representation and the R1 and R5 (the first and the last bead of the planar part) are labeled.

For the MD simulations, we use the coarse-grain MARTINI force field to allow sufficient sampling for free energy calculations. Specifically, we simulated the dimerization of two identical TMDs in an ergosterol containing bilayer environment upon variation of phospholipid composition. We specifically focused on the degree of saturation and presence or absence of anionic phosphatidylserine (PS) in the inner PM leaflet. The chosen phospholipid and ergosterol concentrations and asymmetric distribution mimic key features of the yeast PM. TMD dimer formation is then characterized via the equilibrium ratio of left- and right-handed dimers as well as by the free energy of dimer formation. Both properties are quantified using umbrella sampling. To increase the level of information we then compare results from a large number of independent simulations of TMD dimer formation, starting from two separated TMDs. Finally, we show how the bidirectional interplay between TMDs and lipids can act as a self-regulating process to stabilize dimers. Surprisingly, despite the direct interaction of anionic PS head groups with the TMD in our MD simulations, omission of PS turned out to have no impact on the free energy of TMD dimer formation. This result is consistent with high-resolution live cell imaging experiments, where the network factor of the Slg1^TMD^ in the yeast PM is not affected by the depletion of cellular PS.

## Methods

### Experimental basis

#### Plasmids and yeast strains

The Slg1^TMD^-plasmid RWC1583 (pRS316 backbone, URA3 marker, pPma1-Slg1^TMD^-5xGA-mNeonGreen) was constructed using standard molecular biology techniques to express ANVGAIVGGVVGGVVGAVAIALCILLIVRHIN-GAGAGAGAGA-mNeonGreen from the strong Pma1 promoter. The construct was verified by sequencing. Transformation into yeast cells was performed using the lithium acetate method. The yeast strains created in this study all were derived from the reference strain BY4741 and are listed in Table 1.

**Table 1:** Yeast strains used in this study.

| Strain | Genotype | Source |
| --- | --- | --- |
| BY4741 | MATa his3Δ1 leu2Δ0 met152Δ0 ura32Δ0 | Euroscarf |
| RWS1616 | BY4741 cho1::natNT2 | Spira, 2012 |
| RWS5149 | BY4741 RWC1583 | This study |
| RWS5150 | BY4741 cho1::natNT2 RWC1583 | This study |

#### Yeast cell culture

Plasmid-containing strains were grown overnight in synthetic complete media without uracil supplemented with 2% glucose and 1 mM ethanolamine (SCD - Ura + etn) media at 30*^◦^*C with shaking. For imaging in mid-logarithmic phase, cells were washed 3x with H2O, diluted to OD600 = 0.1 in SCD - Ura + etn and grown for an additional 2.5 h at 30*^◦^*C.

#### Microscopy

Epifluorescence and total internal reflection fluorescence microscopy (TIRFM) were performed on a fully automated iMIC stand (FEI/Till Photonics) with an Olympus 100 x 1.45 NA objective. DPSS lasers (75 mW) at 491 nm (Coherent Sapphire) and 561 nm (Cobolt Jive) were selected through an acousto-optical tunable filter. Images were collected with an Andor iXON DU-897 EMCCD camera controlled by the Live Acquisition (Till Photonics) software. For imaging, coverslips (Knittel Glass No. 1) were cleaned by sonication in absolute ethanol (Sigma), >99.5% acetone (Sigma), 1 M NaOH (Roth), ddH2O and finally stored in absolute ethanol. To immobilize cells, coverslips were coated with 1 mg/ml concanavalin A (Sigma).

#### Image processing and analyses

Microscopy of the yeast PM was performed as described previously.^33^ Images were processed using Fiji and MATLAB (MathWorks Inc., Natick, MA). Images were contrast-adjusted and scaled for presentation purposes only. All TIRFM images were deconvolved using the Lucy — Richardson algorithm in MATLAB using 12 iterations. The required point spread functions were calculated from images of 100 nm tetra spec beads images using the respective excitation and emission wavelengths. The network factor was previously introduced as “intensity distribution” in ref. ^2^ and describes the normalized lateral distribution of yeast proteins at the PM. In essence, the network factor represents the area above cumulative intensity histograms for deconvolved total internal reflection microscopy (TIRFM) images. It represents an intuitive measure for the heterogeneity in the lateral distribution of a specific PM marker. Diffuse arrangements that resemble network-like patterns yield large network factors, whereas strong clustering in patch-like structures results in low values. Calculation of the network factor was performed using a custom-written MATLAB script. All images shown were scaled 3x using Fiji.

#### Determination of lipid composition

Lipid-extractions, mass-spectrometric analyses, quantification, and data processing were performed by Lipotype GmbH (Dresden, Germany). To generate samples, yeast cultures were incubated overnight in SCD + 1 mM choline at 30*^◦^*C with shaking, diluted to an OD600 of 0.1 in SCD + 1mM choline and further grown at 30*^◦^*C to mid-logarithmic phase. 20 OD were harvested for extract preparation and shipped to Lipotype. Lipidome-results in this study were normalized to total lipids of selected categories and compared using Excel. To determine the typical lipid composition of yeast membranes in our culture conditions, we performed lipidomic analysis. For the purpose of our study, we focused our analysis on mixtures of only three of the major lipid constituents in the yeast PM: ergosterol (the main yeast sterol), the apolar phosphatidylcholine (PC) and the anionic PS that make up 13, 16 and 5% of all lipids in yeast membranes, respectively (Figure 2A). Values in Figure 2A and B do not add up to 100% as only three of the 9 lipid classes detected are shown. Note that both ergosterol and PS are strongly concentrated in the yeast PM^34, 35^ and PS is highly restricted to the inner bilayer leaflet. ^34, 36, 37^ The effective concentrations of ergosterol and PS in the PM are therefore far higher than the average values, ^36^ and are close to the chosen concentrations in our simulations (20% ergosterol in each leaflet, 27% PS in the inner leaflet). We also used the Δcho1 mutant that is not able to synthesize PS anymore (Figure 2B). Note that PC and PS in yeast slightly differ in their degree of saturation (Figure 2C) and acyl chain lengths (Figure 2D), with most PC containing two unsaturated and shorter acyl chains (C16:1/C16:1, C16:1/C18:1) and PS featuring more mixed chains (C16:0/C18:1, C16:1/C18:1, C16:1/C18:0). Also note that budding yeast only expresses a single desaturase enzyme so that acyl chains typically contain no more than a single double bond.

**Figure 2:**
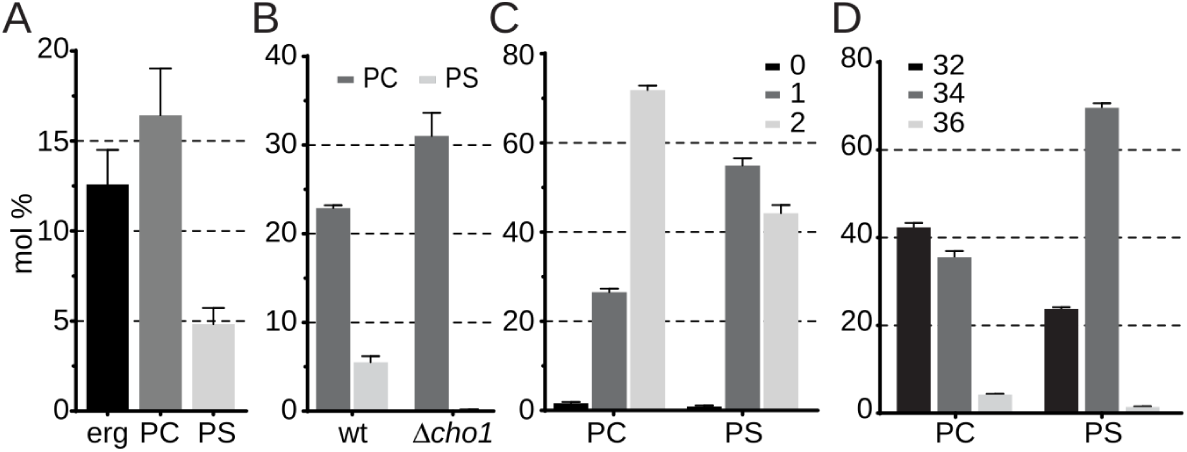
Lipid composition. Lipid composition of yeast membranes. (A) The relative amounts of ergosterol, PC and PS in control cells at 30*^◦^*C, (B) The depletion of PS in the Δcho1 mutant, (C) the acyl chain saturation for PC and PS and (D) the length of acyl chains for PC and PS are represented.

### Simulation basis

#### Choice of the model

The initial atomistic structure of the Slg1 TMD was prepared via SWISS-MODEL homology modeling webserver. ^38^ The structure was replicated in a way that the two monomers are located at the distance of 4.5 Å from each other. The two monomers were inserted in membranes with different compositions using Martini maker as implemented in web-based CHARMM-GUI membrane builder. ^39^ Accordingly, the systems were solvated by water molecules and ions were added to neutralize the systems. The final systems consist of two TMD domains and around 400 lipid molecules (Figure 1B). Five different systems with different compositions were prepared. All the systems contain 20% ergosterol. One system with POPC (16:0,18:1) and ergosterol (POPC-Erg) was additionally created. Two systems out of five have the most similar lipid composition as the implemented experiments (previous section), one without PS lipids (DOPC-POPC-Erg) and the other with PS lipids (DOPC-POPC-Erg-PS). As noted in the previous section, a realistic representation of the yeast PM would indeed include all major phospholipid constituents (PC, PE, PI, PS), ergosterol as well as complex sphingolipids in the extracellular leaflet. However, an adequate Martini representation of the most prevalent yeast sphingolipids, Mannosyl-inositolphosphoryl-ceramides, is still lacking. These sphingolipids are thought to contribute to gel-like PM microdomains in yeast, which were not the aim of our current study. Our goal is therefore not to completely replicate the yeast PM but only maintain some of its key features (mainly of the inner leaflet) and to obtain mechanistic insights on weak TMD-lipid interactions, which requires the simulations to approach steady state and this can be achieved by limiting the complexity of our lipid composition. Indeed, we focused on PC variants to better relate to the large number of existing studies on the interaction of this phospholipid with TMDs. We then included ergosterol as this sterol was expected to directly interact with the GxxxG motifs of our chosen TMD. Finally we included PS on the inner leaflet. This lipid makes up the major portion of charged lipids on the inner leaflet of the PM and this negative charge is an important characteristic of the PM interface to the cytosol. The amount of PS lipids in our model membranes is 27% of lipids in the lower leaflet. It should be noted that due to the coarse-grain structure of the lipids in MARTINI, not all the lipid types in the actual experiment could be reproduced also with respect to the saturation status of the chains, and therefore, the lipid composition is in general only an approximation. Two extra systems were also constructed similar to the experimental composition with the exception that DOPC (18:1,18:1) is replaced by a more unsaturated DIPC (18:2,18:2) lipid (DIPC-POPC-Erg, DIPC-POPC-Erg-PS) in order to probe the effect of saturation status. The percentage of each lipid type in all membrane systems is represented in Table 2.

**Table 2:** Percentage of the lipid components for each system, averaged over each leaflet. Note that the PS lipids are present only in the lower leaflet.

| **System** | leaflet | POPC | DOPC | DIPC | POPS | DOPS | DIPS | Erg |
| --- | --- | --- | --- | --- | --- | --- | --- | --- |
| POPC-Erg | upper | 80 | 0 | 0 | 0 | 0 | 0 | 20 |
|  | lower | 80 | 0 | 0 | 0 | 0 | 0 | 20 |
| DOPC-POPC-Erg | upper | 26.7 | 53.3 | 0 | 0 | 0 | 0 | 20 |
|  | lower | 26.7 | 53.3 | 0 | 0 | 0 | 0 | 20 |
| DOPC-POPC-Erg-PS | upper | 26.7 | 53.3 | 0 | 0 | 0 | 0 | 20 |
|  | lower | 15.2 | 38.1 | 0 | 7.6 | 19.1 | 0 | 20 |
| DIPC-POPC-Erg | upper | 26.7 | 0 | 53.3 | 0 | 0 | 0 | 20 |
|  | lower | 26.7 | 0 | 53.3 | 0 | 0 | 0 | 20 |
| DIPC-POPC-Erg-PS | upper | 26.7 | 53.3 | 0 | 0 | 0 | 0 | 20 |
|  | lower | 15.2 | 38.1 | 0 | 7.6 | 19.1 | 0 | 20 |

Since two systems out of all simulated systems include PS lipids in the lower leaflet and as a result are asymmetric, we have checked the pressure profiles, which are symmetric across the bilayer (Figure S1).

#### Simulations of TMD association

To study the association of the Slg1 TMDs in different bilayer systems, coarse-grain molecular dynamics (MD) simulations were performed. To describe the interactions between the peptides, lipids and solvent molecules, the version 2.2 of MARTINI force field was used. ^40–42^ In MARTINI force field, on average 4 atoms along the associated hydrogens are represented by one interaction site (bead). We would like to mention that the reason we have not performed the newest version of MARTINI, i.e., version 3.0 is that the parameters for sterol molecules are not available yet and in our model systems the role of ergosterol is crucial.

The MD simulations were performed using the version 2019 of GROMACS.^43, 44^ The constant temperature was controlled by coupling the system to a Berendsen thermostat ^45^ at 310 K. The pressure was controlled by coupling the system to a barostat (coupling time of 0.1 ps and compressibility of 3 *×* 10*^−^*^4^ bar*^−^*^1^) at 1 bar, using semi-isotropic coupling scheme. ^45^ For the production simulations the constant pressure was controlled using the Parinello-Rahman barostat. ^46^ The non-bonded interactions were treated using a switch function from 0.0 to 1.1 nm for the electrostatic, and from 0.9 to 1.1 nm for Lennard Jones interactions. For the DOPC-POPC-Erg-PS system we additionally set up simulations using the polarizable water model of MARTINI force field. ^47^ The equilibration procedure was performed according to the input files provided by the CHARMM-GUI webserver. For each system, 29 independent samples were simulated, starting from the last structure from the equilibration process by reinitializing velocities. Since the same starting structure has been used for each sample, we checked if there is a bias related to this structure by calculating the autocorrelation function of the temporal change of the angle between the two TMDs (Figure S2). Accordingly, the correlation time for the DOPC-POPC-Erg system is 32 ns, whereas the average latest time at which the two TMD are at a distance of 3 nm (the distance at which the two TMD start to interact with each other) is around 2 *µ*s for this system. This latest time can also been observed for one sample system (Figure S3). Therefore, the correlations times are far lower than this time and no bias can be expected. The simulation time for each sample is 5 *µ*s performed in the NPT ensemble with the time step of 20 fs. In the rest of the paper, we refer to these simulations as ‘ensemble’ simulations.

All the simulations data have been analyzed using the python routines incorporating the MDAnalysis package^48, 49^ and GROMACS tools. The VMD software was used to visualize the trajectories. ^50^

#### Crossing angle and tilt angle of TMDs

The crossing angle was calculated based on ref. ^51^ Accordingly, the crossing angle is the dihedral angle constructed by the beginning of each helix and the two points on the two helices, which are at the closest distance from each other. The beginning of the helix is considered as the center of mass (COM) distance of the backbone of the last three residues on the N-terminus side of the helices. The right-handed (RH) and left-handed (LH) configuration correspond to negative and positive crossing angles, respectively.

The average tilt angle of the helices was calculated as the angle between the vector connecting the beginning and end of each helix and the membrane normal (z-axis). The beginning and ending of the helix was considered respectively as the COM of the first three and last three residues of the helix, considering the backbone beads of the proteins.

#### Potential of mean force

In order to estimate the stability of the dimer in different membranes, we calculated the potential of mean force (PMF) profile for the TMDs dissociation. For this purpose, we used umbrella sampling (US).^52^ The reaction coordinate was chosen as the COM distance of the two TMDs. For the simulations starting from a bound state, the starting structurein which a RH or LH dimer is formed was extracted from ensemble simulations, and then one TMD was pulled using the pull scheme of GROMACS with the rate of 0.1 nm per ns, while the other TMD is restrained with a force constant of 1000 kJ mol*^−^*^1^ nm*^−^*^2^. In each window, the umbrella potential was applied so as the COM distance is restrained with the force constant of 1000 kJ mol*^−^*^1^ nm*^−^*^2^. 29 windows were run with the window size of 0.1 nm and the reaction coordinate was restrained with a force constant of 1000 kJ mol*^−^*^1^ nm*^−^*^2^. Each window was first equilibrated for 50 ns, followed by 8 to 12 *µ*s production simulations. The WHAM method was employed to unbias the umbrella potentials and to combine all the windows. ^53^ For the final energy profiles, we estimated the standard deviation by defining four independent non-overlapping fragments of each MD simulation. To check any hysteresis effect due to the definition of reaction coordinate, two simulations for the DOPC complex mixtures were performed and for these simulations, one starting structure was chosen from US simulations of the bound state in which the distance of the two TMDs is around 3 nm and then for the subsequent windows the equilibrated structure (with equilibration time of 50 ns) of the previous windows was used as a starting structure until the whole range of reaction coordinate is covered.

#### Membrane thickness, order parameter of phospholipids and tilt angle of ergosterol

The membrane thickness was estimated by calculating the density of the head groups of phospholipids, using the density tool of GROMACS. The membrane thickness was subsequently defined as the distance between the two maximum values of the density profile, corresponding to the upper and lower leaflet of the membrane.

The order parameter measures how the lipid chains are oriented with respect to the membrane normal and quantifies the degree of their orientational order. According to ref. ^54^ the order of lipid chains can be quantified using molecular order parameter (*S_mol_*)

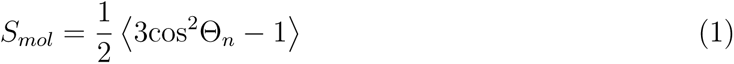

where Θ*_n_* is the angle between the vector constructed by n^th^ segment of the hydrocarbon chain, i.e., *C_n−_*_1_ and *C_n_*_+1_, connecting the *n −* 1 and *n* + 1 carbon atoms, and the membrane normal (z-axis). For the coarse-grain lipids, however, the vector connecting the two consecutive beads is considered, i.e., *C_n−_*_1_ and *C_n_*. Subsequently, this quantity is calculated for all the beads of the two chains separately. The angular brackets represent the time and ensemble average.

The tilt angle of ergosterol was analyzed considering the planar part and was taken as the angle between the vector connecting R1 and R5 beads and the membrane normal (z-axis) (Figure 1B).

## Simulation results

To systematically study the interactions between a yeast TMD and the lipid bilayer, we simulated the dimerization/association of two Slg1 TMD molecules (Figure 1A), using coarse-grain MD simulations (Figure 1B), and explored the effects of lipid environment on the TMD association — and vice versa.

### Dimerization properties induced by membrane environment

#### Association of TMDs

The self-association of the TMDs was studied. In the initial structure, the two TMDs were separated with the distance of 4.5 nm in order to eliminate the initial interactions between monomers (Figure 1B). During the 5 *µ*s simulation for each sample of ensemble simulations (see Methods section), on average the dimer is formed in 92.8% of all the independent simulation samples, considering all systems; see Figure S3 for a representative example.

Before the dimer is formed, a particular pathway is followed. Here, the tilt angle can describe how the two TMDs approach each other until a compact dimer is formed (Figure 3). Indeed, the average tilt angle of the TMDs is reduced upon dimerization and a systematic distance dependence is observed: as the two TMDs approach each other, the tilt angle is reduced. For 12<COM<*∼*20 Å, it continuously decreases as the two TMDs come closer to each other. This region may be interpreted as a pre-dimerized state, in which the two TMDs interact mainly through the C-terminus residues, in a way that a “V”-shaped structure is formed. This is also reflected in the average number of contacts between the corresponding similar residues on the two TMDs for 10<COM<20 Å(Figure S4). For COM<8 Å, the tilt angle is the lowest and is nearly constant (dimer). Here we choose 10 Å to define the transition between the dimer state and pre-dimerized state.

**Figure 3:**
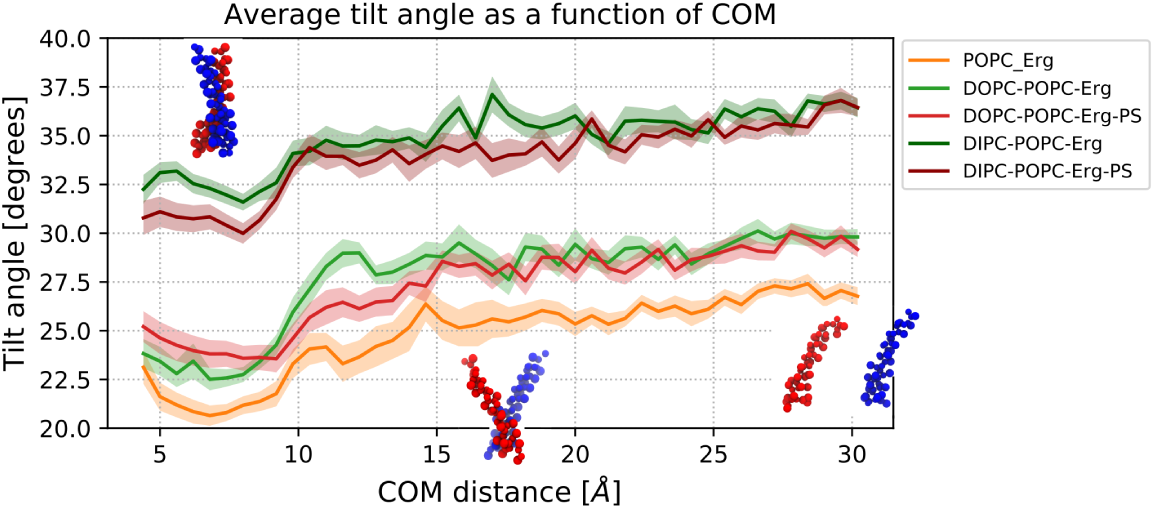
The of the two TMDs as a function of the COM distance between the two TMDs. The snapshots of the two TMDs are also shown for different COM distances. The shaded area represents the standard error of the mean determined using ensemble simulations.

The tilt angle of TMDs strongly depends on the chosen lipids. In DOPC systems the tilt angle is significantly higher than in DIPC systems, whereas the presence of the PS lipids has a negligible impact.

The configuration of the two TMDs with respect to each other and the final dimer structure can be characterized by a crossing angle (see simulation basis). Accordingly, there are two possible configurations for both the pre-dimerized state and final dimer: right-handed (RH) and left-handed (LH) crossing angles. At large COM distances, where the TMDs are far from each other, the crossing angle is around zero due to symmetry reasons (Figure 4A). As the two TMDs approach each other, the average crossing angle increases and, interestingly, it displays a similar behavior for all the systems before they start to enter a dimerized state, mainly for the range 15<COM<20 Å(Figure 4A,B), since it is mainly positive, and for the systems with higher saturation in lipid chains. This means that the LH configurations are the main intermediate structures, which can subsequently lead to either RH or LH final dimers. This positive average value is a consequence of a broad distribution of the positive values (reaching values up to 75*^◦^*) as compared to the distribution of negative values (Figure 4B). Generally speaking, this suggests that there is nearly a universal path towards the formation of the binding state. The binding state itself displays a much stronger dependence to the lipid composition, discussed as follows.

**Figure 4:**
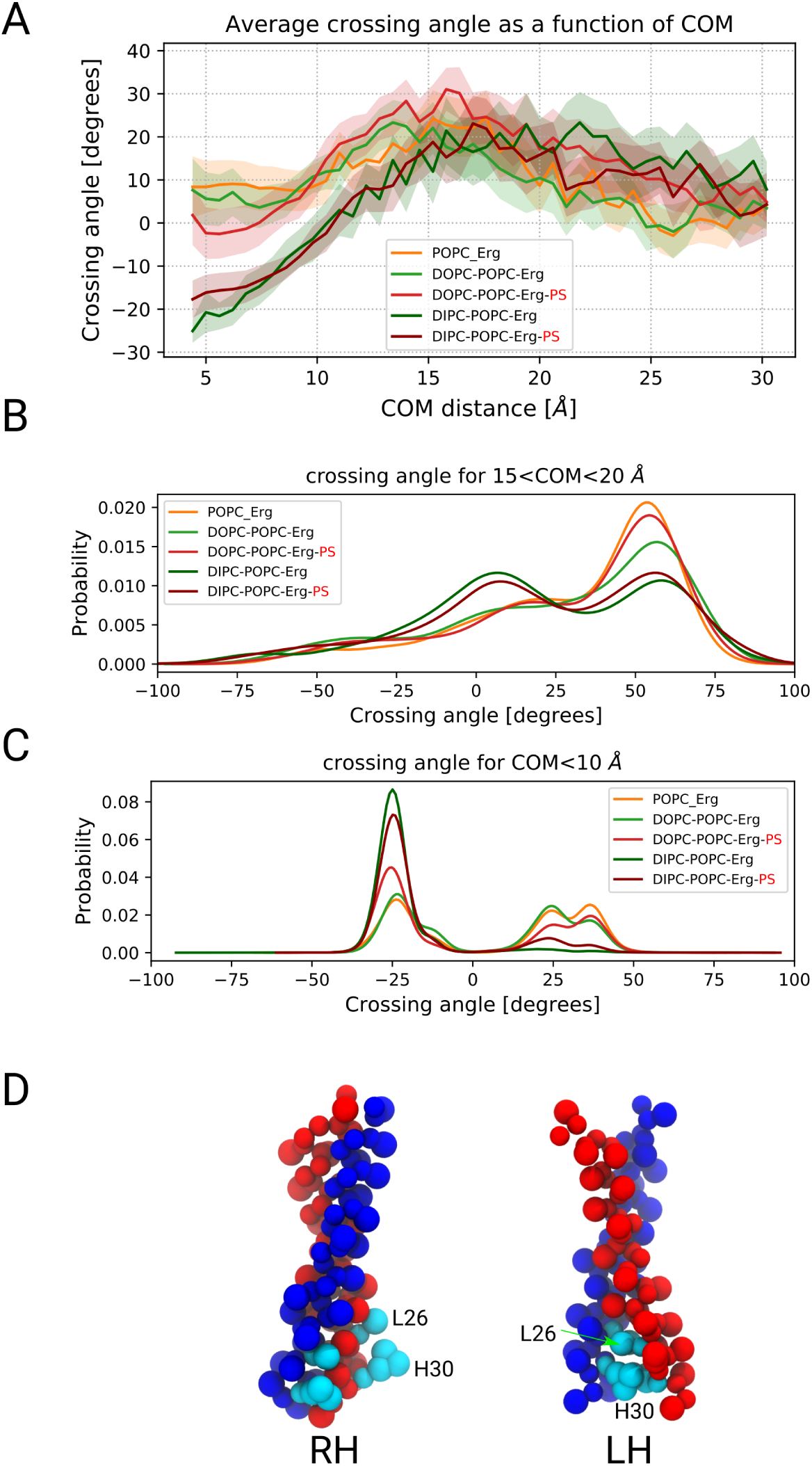
(A) The average crossing angle of the two TMDs as a function of COM distance between the two TMDs is shown. (B) The distribution of crossing angle for 0<COM<10 Å and (C) for 10<COM<20 Å is shown. (D) The RH and LH configurations of the dimer are shown. The L26 and H30 residues are depicted in cyan.

Upon dimerization in ensemble simulations, the crossing angle values are decreased and the form of the distribution depends on the system (Figure 4C). The complex mixtures with DIPC lipids represent mainly RH configurations, whereas POPC-Erg as well as complex mixtures with DOPC lipids form both RH and LH dimers, with nearly similar probabilities (Table 3). Interestingly, LH structures are less well-defined than RH structures, as reflected by their broader distribution of crossing angles: the average crossing angle for RH configurations is *∼*25*^◦^*, whereas LH configurations exhibit either similar or higher values of crossing angle when compared to RH ones. In LH configurations, L26 and H30 residues, which have relatively longer and bulkier side chains, face each other, while in RH configurations they are relatively far from one another (Figure 4D). As noted above, these residues have a significant role also in pre-dimerization via the C-terminus and their interaction initializes the dimerization process (Figure S4). Therefore, the universal pathway in the pre-dimerized state, as discussed above, is expected since the attachment of these residues induces LH configurations.

**Table 3:** The percentage of RH and LH configurations collected from all the association samples in ensemble simulations in which the dimer is formed as well as the ones extracted from the associated ΔG values is shown.

| **System** | **%RH in ensemble sim.** | **%RH estimated using $\Delta G$** |
| --- | --- | --- |
| POPC-Erg | $48 \pm 9$ | 73.0 |
| DOPC-POPC-Erg | $54 \pm 9$ | 53.7 |
| DOPC-POPC-Erg-PS | $56 \pm 9$ | 23.5 |
| DIPC-POPC-Erg | $86 \pm 6$ | - |
| DIPC-POPC-Erg-PS | $79 \pm 7$ | - |

Furthermore, it should be also mentioned that the simulation of the dimers via atomistic CHARMM-based simulations is an ongoing project in our group. As a preliminary result, to be shown in the SI (Figure S5), we can already report that the dimer structures of the averaged MARTINI model are very similar to the back-mapped atomistic system. Furthermore, after mapping the CG system to the atomistic system the dimer is stable throughout the microsecond timescale simulations. Thus, we may tentatively conclude that there are no serious artifacts in the MARTINI model.

#### Residue contacts

Next we looked at the TMDs interfaces in the dimer state in ensemble simulations, specifically for RH and LH configurations. For this purpose, we calculated the contact map of the TMDs residues for 0<COM<10 Å (Figure 5). For POPC-Erg and DOPC systems, the number of residue contacts for RH and LH state is rather similar, which is in line with the similar probabilities of RH and LH configurations (Table 3). For DIPC systems, however, we observed a significant higher number of residue contacts for RH and the opposite for LH configurations (Figure 5). This is in agreement with the dominating probability of RH configurations in these systems. If we count the number of contacts for the range where the COM distance between each similar residue pair on the two TMD is between 0 and 9 Å, this number, for instance, for DOPC systems is 29 and 24, for RH and LH configurations, respectively, whereas the corresponding numbers for DIPC systems are 52 and 8. This shows the similarity of the number of contacts for RH and LH configurations in DOPC system and a considerable difference in DIPC systems.

**Figure 5:**
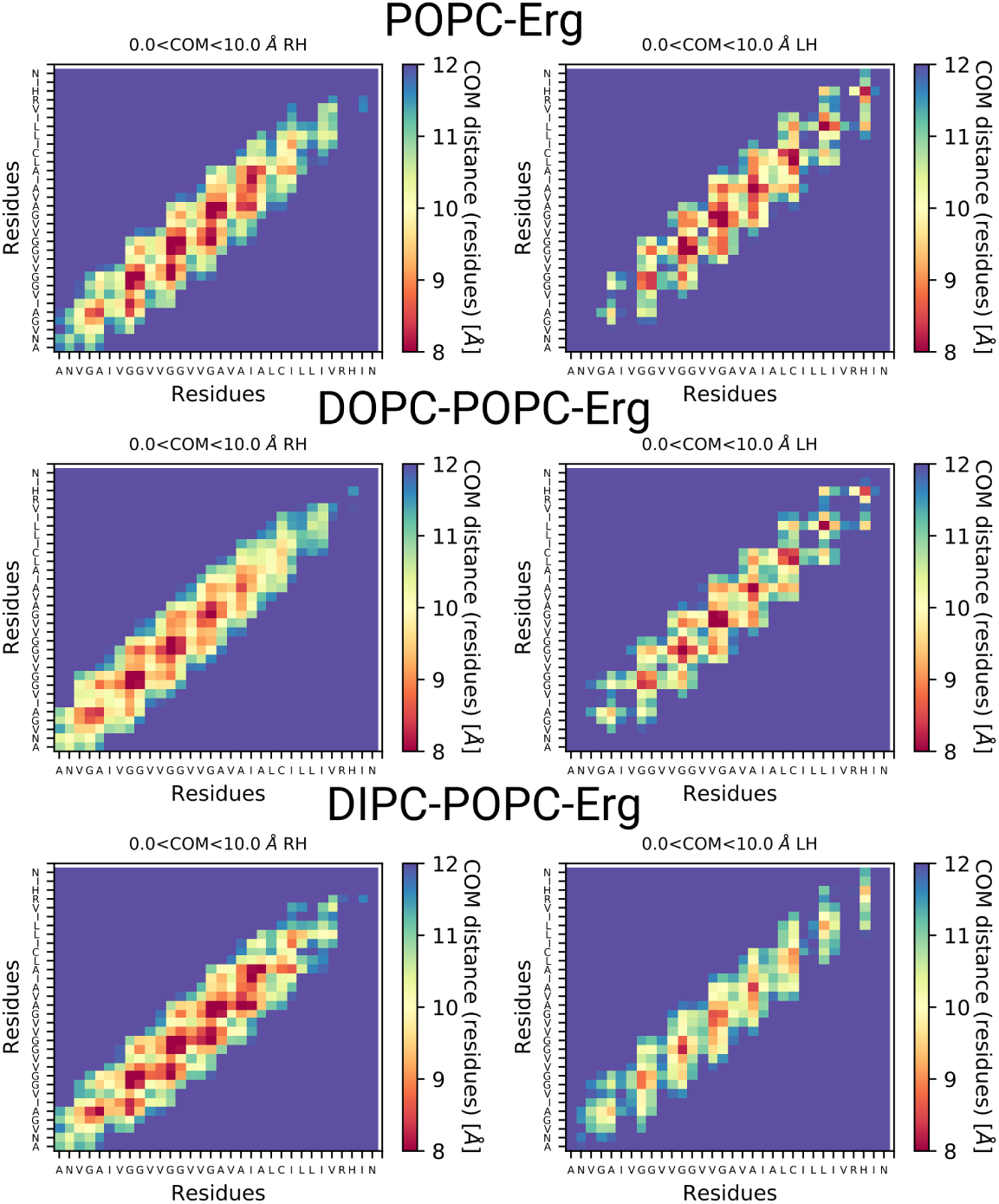
The contact maps of the TMD residues in different membrane systems for RH (left) and LH (right) configurations is shown. The COM distance between each residue pair is considered as a contact.

Furthermore, considering different parts of the dimer, in DOPC systems, a considerable binding is taking place also via C-terminus residues, which is mainly due to a significant number of LH configuration, as discussed earlier. In DIPC systems, however, the binding is established mainly via the N-terminus and central residues of the TMDs and the role of C-terminus residues is markedly lower, even in LH configurations. This is probably due to the higher tilt of the TMDs in the DIPC systems and the dominating number of RH configurations.

#### Energetics of dimerization

In order to estimate the stability of the dimer in different lipid environments, we quantified the free energy of dissociation, i.e., the PMF as a function of the COM distance of the two TMDs via umbrella sampling (see Methods) and the energy was converged after 6 to 9 *µ*s (Figure S6). The shape of the free energy profiles represents a global minimum and the position of the minimum is around 0.7 nm (Figure 6A) for all systems. As it was observed earlier, POPC-Erg and DOPC systems ended up in both RH and LH configurations with nearly similar probabilities. Therefore, for these systems we performed the PMF calculations both for RH and LH configurations. For the RH configurations we performed PMF calculations starting from both bound and unbound states to make sure that similar results are obtained (Figure S7). Now to check if the final structure of the bound and unbound simulations are similar, we have calculated the crossing angle distribution for the window in which the COM distance is restrained at 0.7 nm, corresponding to the minimum of the free energy curve, which show very similar values for the bound and unbound simulations (Figure S8).

**Figure 6:**
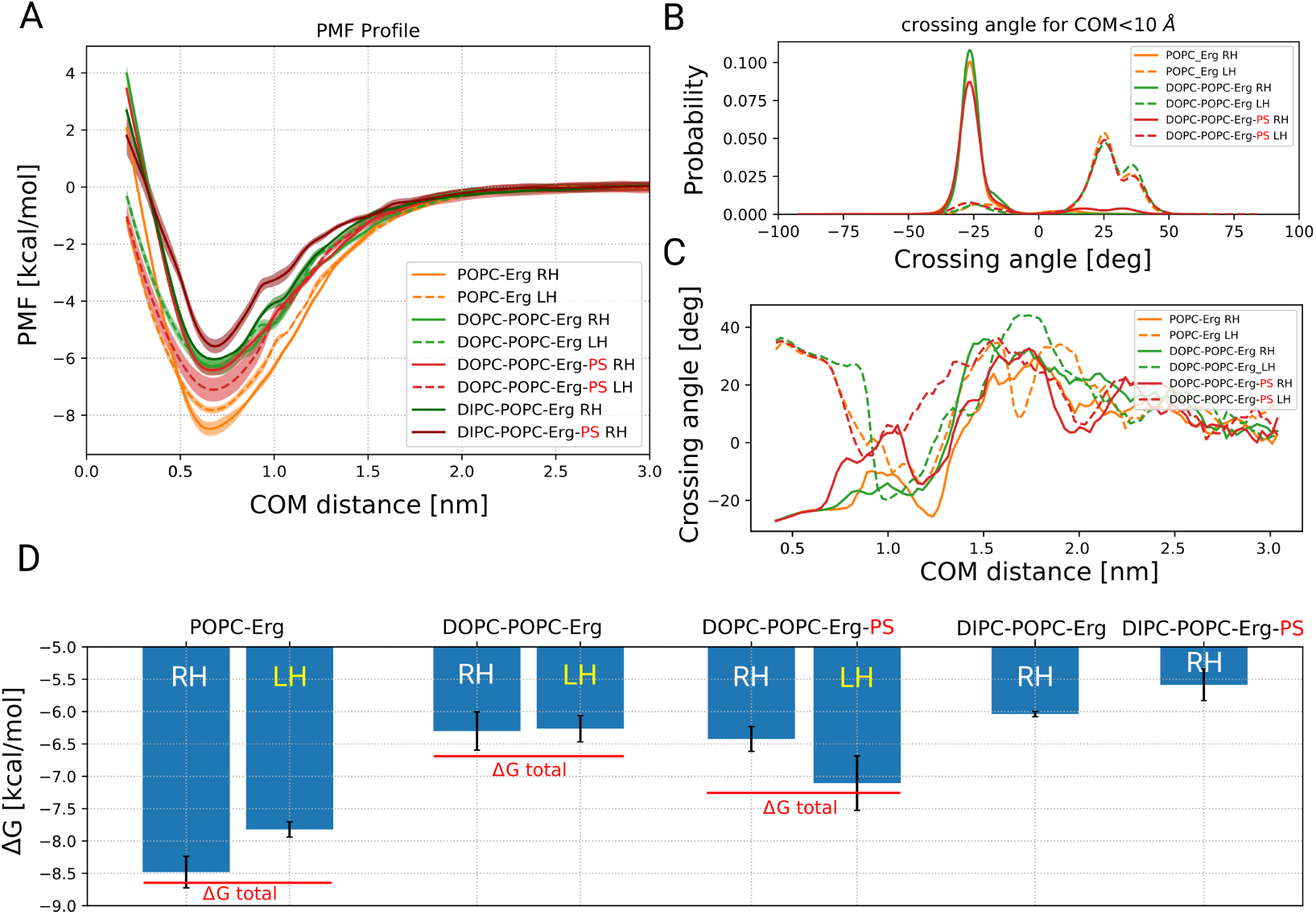
(A) The free energy profiles of the dimerization for all systems for RH starting structure is represented. For POPC-Erg and DOPC complex mixtures, the corresponding profiles for LH initial structure is also represented. (B) The distribution of the crossing angle and (C) the variation of crossing angle as a function of COM distance between the two TMDs in the US simulations are shown. (D) The corresponding ΔG values, i.e., the minimum of the free energy profiles in (A) are depicted. The shaded areas and the error bars show the uncertainty, which are explained in Methods.

For all free energy calculations, the crossing angle values remain either RH or LH and do not switch from one to another (Figure 6B). Interestingly, a similar behavior is observed in the dimerization pathway as the ensemble simulations: irrespective of the initial configuration, the crossing angle is mainly LH in the pre-dimerized state (Figure 6C). Also interestingly, LH configurations represent weaker helix-helix interactions as compared to the RH ones in POPC-Erg and similarly in DOPC systems without PS lipids (Figure 6A,D). The opposite behavior as for the POPC-Erg system is observed in DOPC complex mixtures with PS lipids. This might be due to the fact that in the LH configurations, the charged residues are facing each other (Figure 4D) and are hardly accessible by the PS lipids, and probably, reducing the protein-lipid interaction and stabilizing protein-protein interaction. These differences are, however, not dramatic. Considering these two different states, i.e., RH and LH, we can estimate the total free energy of dimerization: 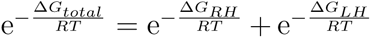, where R is the gas constant and T is the room temperature. Accordingly, stronger helix-helix interaction is predicted for the POPC-Erg and DOPC systems (Figure 6D). Subsequently, we estimated the population of RH and LH configurations based on the free energy profiles. It is possible to obtain these populations using the corresponding ΔG values (Table 3). Alternatively, a more precise way would be to calculate the integral of the corresponding free energy profiles over the entire reaction coordinate and from that to extract the probabilities. However, this integration over the phase space hardly changed the compositions. The probabilities for RH and LH configurations obtained from the free energy of dimerization are different from the ensemble simulations. For the POPC-Erg system, the free energy predict higher probabilities for RH configurations when compared to the ensemble simulations (Table 3). The results are also opposite for DOPC systems with PS lipids and the free energy predicts a considerably lower number of LH configurations as compared to the ensemble simulations (Table 3). The DOPC system without PS lipids predicted, however, similar probabilities in both approaches. Strictly speaking, since the the ensemble simulations are not in equilibrium and there are no transitions between pre-dimerized and dimerized states, the probabilities of the RH and LH conformers might not agree with the ones estimated from free energy calculations, which are indeed in equilibrium. This likely depends on the nature of the energetic barrier between these states and may explain why for POPC-Erg system, ΔG values suggest a majority of RH configurations, and for DOPC systems with PS, the majority of LH configurations, in contrast to the result of the ensemble simulations.

The final free energies of dimerization for all systems show that the POPC-Erg system represents the strongest helix-helix interaction, whereas the DIPC systems display the weakest interactions (Figure 6D). Therefore, in general, the larger the number of unsaturated bonds of the surrounding phospholipids (PLs), the weaker the helix-helix interaction (DIPC<DOPC<POPC). Furthermore, the presence of PS lipids slightly destabilizes the dimer in DIPC complex mixtures, whereas they have the opposite effect, i.e., they stabilize the dimer in DOPC systems.

#### Dimerization and relation to membrane properties

In this section, we inspect some general properties of the studied membranes. First, we see how the thickness is changed and whether it can be related to the tilt angle of TMDs and the energy of dimerization. We also intend to probe the details of interaction of each lipid type with the TMD, and finally, how the formation of the dimer can affect the properties of the membrane, specifically the order parameter of lipids and the orientation of ergosterol molecules.

#### Membrane thickness

Different lipid molecules have different chain length and saturation state, and therefore, the thickness of the membrane varies in various lipid environments. On the other hand, the TMDs themselves have a specific hydrophobic length, and thus, the TMDs and membrane naturally adjust themselves to each other.^17, 55, 56^ This adjustment requires that the TMDs are tilted if their hydrophobic length is higher than the membrane hydrophobic thickness, which is the case for all the studied membranes in this work. The POPC-Erg system displays the highest thickness and the complex mixtures have nearly similar thickness values (Figure 7). The addition of PS lipids in both complex mixtures can only negligibly change the membrane thickness. Furthermore, the thickness is correlated with the TMD tilt angle, however, small changes in membrane thickness significantly alters the tilt angle of the TMDs. This might mean that thickness is not the only criterion to determine the tilt angle but other aspects of the specific interaction of the protein and the membrane are also involved. The thickness is also slightly correlated with the free energy of association: the membrane with highest thickness, i.e., POPC-Erg represents the strongest helix-helix interaction (Figure 6D).

**Figure 7:**
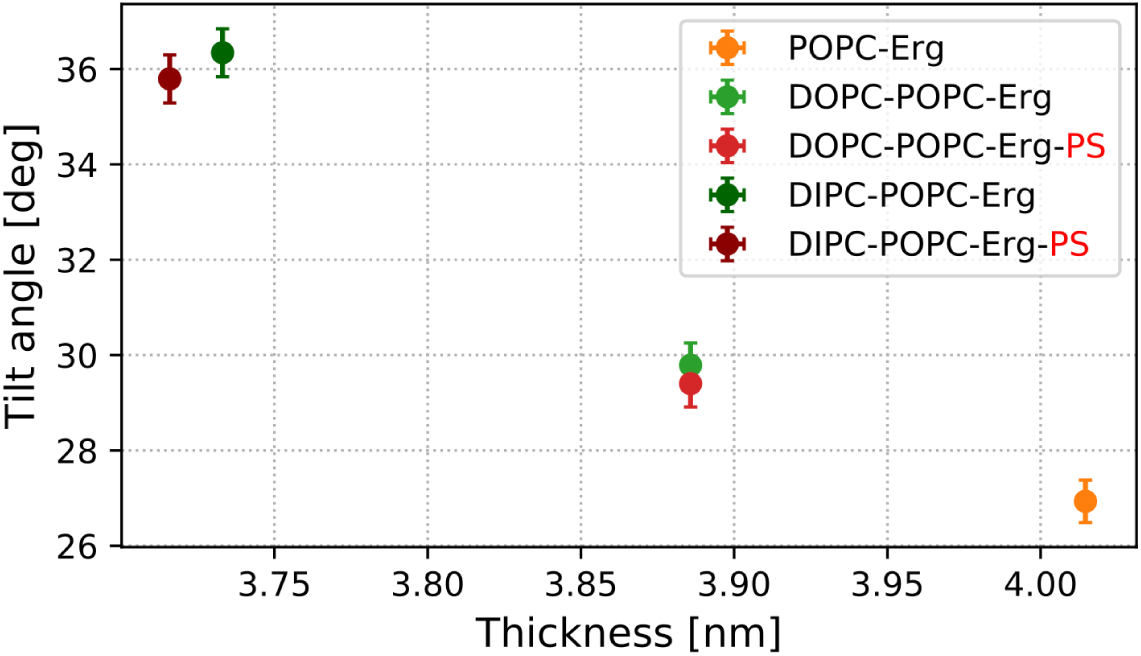
Average tilt angle of TMDs as a function of membrane thickness. The error bars are the standard error of the mean determined using ensemble simulations.

#### Radial distribution function

In order to probe the lipids packing around the dimer in complex mixtures in the ensemble simulations, we quantified the radial distribution function (RDF) of the COM of the two TMDs and the head group of each lipid type in the membrane (Figure 8A,B). In all systems, the RDF profiles show very strong interaction of ergosterol with the TMD. Interestingly, this interaction is stronger with the upper part of the protein, where the GxxxG patterns reside, showing that ergosterol has a strong tendency to interact with these patterns. For the systems containing PS lipids, PS lipids represent a (weak) attractive interaction with the TMDs and approach the dimer quite closely, but not as close as ergosterol. There is a weak reduction of non-PS lipids when PS is added, which is naturally due to the fact that PS lipids take some space. The weak destabilization upon the addition of PS lipids, observed from the thermodynamics (ΔG), is reflected by the presence of weak interactions with these components. Also interestingly, POPC displays an attractive interaction (i.e., g(r)>1) only for DIPC but not for DOPC. Comparing the two complex mixtures, protein-lipid coupling already starts for distances <3 nm for DIPC and <1.7 nm for DOPC. This might be a consequence of the more disorderd nature of DIPC lipids with a correspondingly larger interaction range.

**Figure 8:**
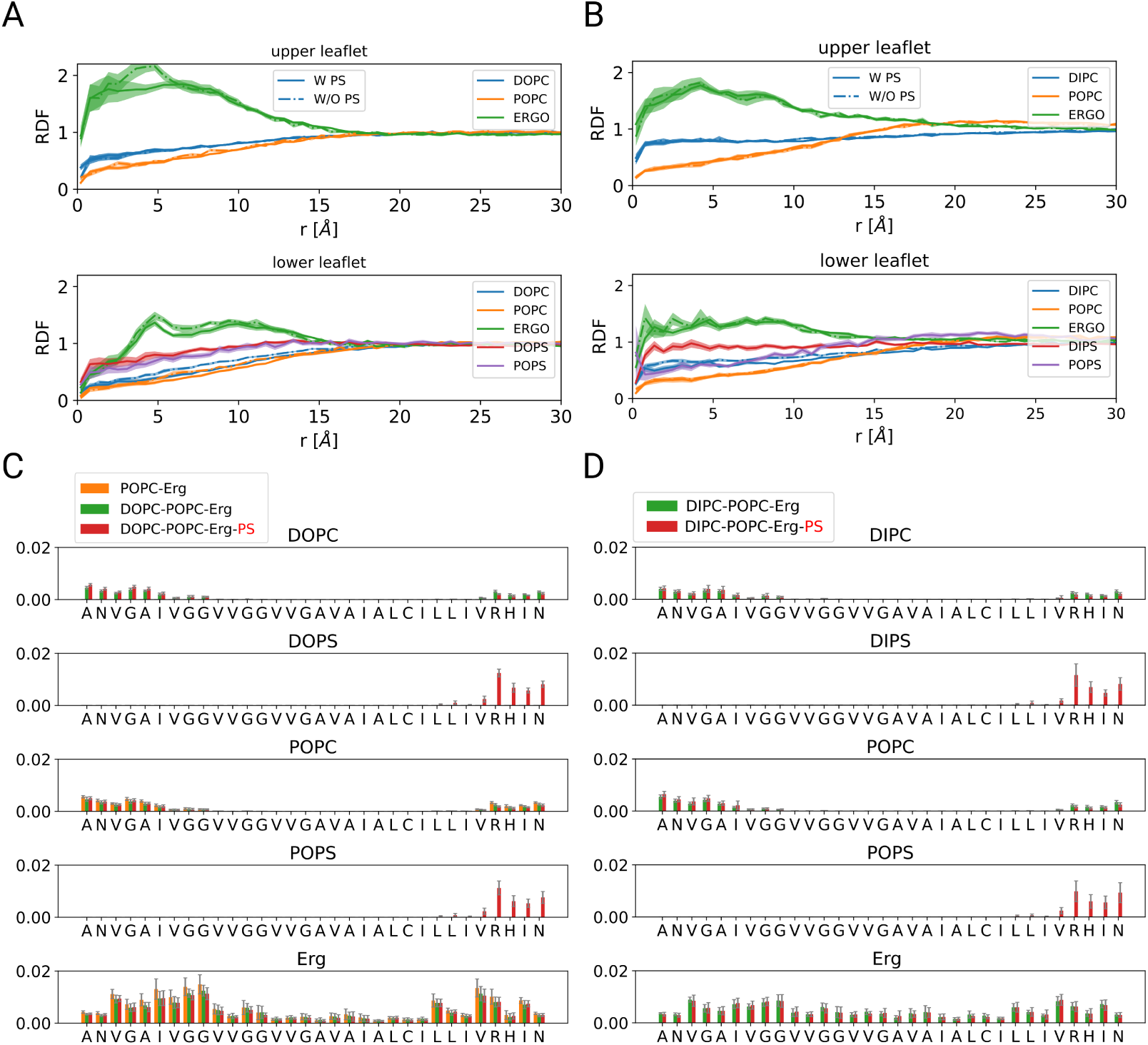
(A) The average radial distribution function of the COM of the two TMDs and the head groups of the lipids is shown for DOPC and (B) DIPC complex mixtures, considering the configurations in which the dimer is formed. (C) The average number of contacts of the lipids head groups with the residues of the TMDs divided by the number of lipids for each type, calculated for the configurations in the dimer state for the POPC-Erg and DOPC and (D) DIPC systems is represented. A contact between each lipid type and a TMD residue is considered to be formed when the distance between the lipid head group and the COM of the considered residue is less than 6 Å. The number of contacts for the two TMDs were subsequently added up. The shaded areas and error bars are the standard error of the mean determined using ensemble simulations.

Furthermore, we performed one set of ensemble simulations using the polarizable water model of MARTINI force field and calculated the RDF profiles (Figure S9). The results show a little bit higher interaction of PS lipids with the dimer at close distance. However, the order of the interaction as well as its limit at large distances is very similar compared to the conventional MARTINI model.

#### Lipid-TMD contacts

The RDF profiles of the TMDs and lipids, discussed in the previous section, provide a macroscopic picture of the lipid-protein interaction and does not give a detailed description of the interaction of the lipids with the TMD residues. Therefore, here we have analyzed the contacts of each lipid molecule with all the TMD residues in the dimer state for the POPC- Erg and the complex mixtures in ensemble simulations (Figure 8C,D). In agreement with RDF profiles, ergosterol interacts significantly with the TMDs, both in the intracellular and the extracellular side, especially with the GxxxG patterns. Interestingly, the interaction of ergosterol with the TMDs in DIPC systems is considerably different, representing significant interaction with the amino acids in the middle of the protein (Figure 8D). This might be due to the higher tilt angle of ergosterol in these systems, especially in the proximity of dimer (Figure S10). Moreover, the contact of ergosterol with the TMDs is not noticeably changed by the presence of PS lipids. PS lipids interact significantly with the dimer, mainly with R29, and mostly take the position of DOPC, DIPC and POPC, reflected also in the RDF profile (Figure 8C,D). Moreover, in RDF profiles, POPC displayed an attractive interaction (i.e., g(r)>1) only for DIPC but not for DOPC, which is reflected here by the increased size of the bars. However, this also implies that the absolute values of the number of contacts are somewhat difficult to interpret since there are also contacts although the dimer is exclusively repulsive, i.e., g(r)<1 for all r values.

#### Order parameter of phospholipids

So far, we have observed how the membrane environment affects the protein-protein and protein-lipid interactions. In this section, we want to explore how the formation of the dimer can affect the membrane properties in ensemble simulations. For this purpose, we have quantified the average order parameter of the PLs, which is extensively used to describe the structural properties of the lipid bilayers. Accordingly, we estimated the order parameter both for the monomer (Figure 9A) and the dimer state (Figure 9B). The average lipid order parameter hardly depends on the distance to the protein if the protein is not dimerized (Figure 9A). This means that a monomer does not have an impact on the average behavior of the acyl chains. In striking contrast, there is a much stronger perturbation of the order parameter induced by the formation of dimer (Figure 9B). Indeed, starting from large distances, the order parameter first increases until *∼*1 nm (ordering effect of the dimer). For smaller distances, where all PLs experience a repulsive force, the order parameter is decreased. The increase of order parameter at intermediate distances is systematically higher if the absolute value of the order parameter is higher, i.e., for more saturated systems (Figure 9B).

**Figure 9:**
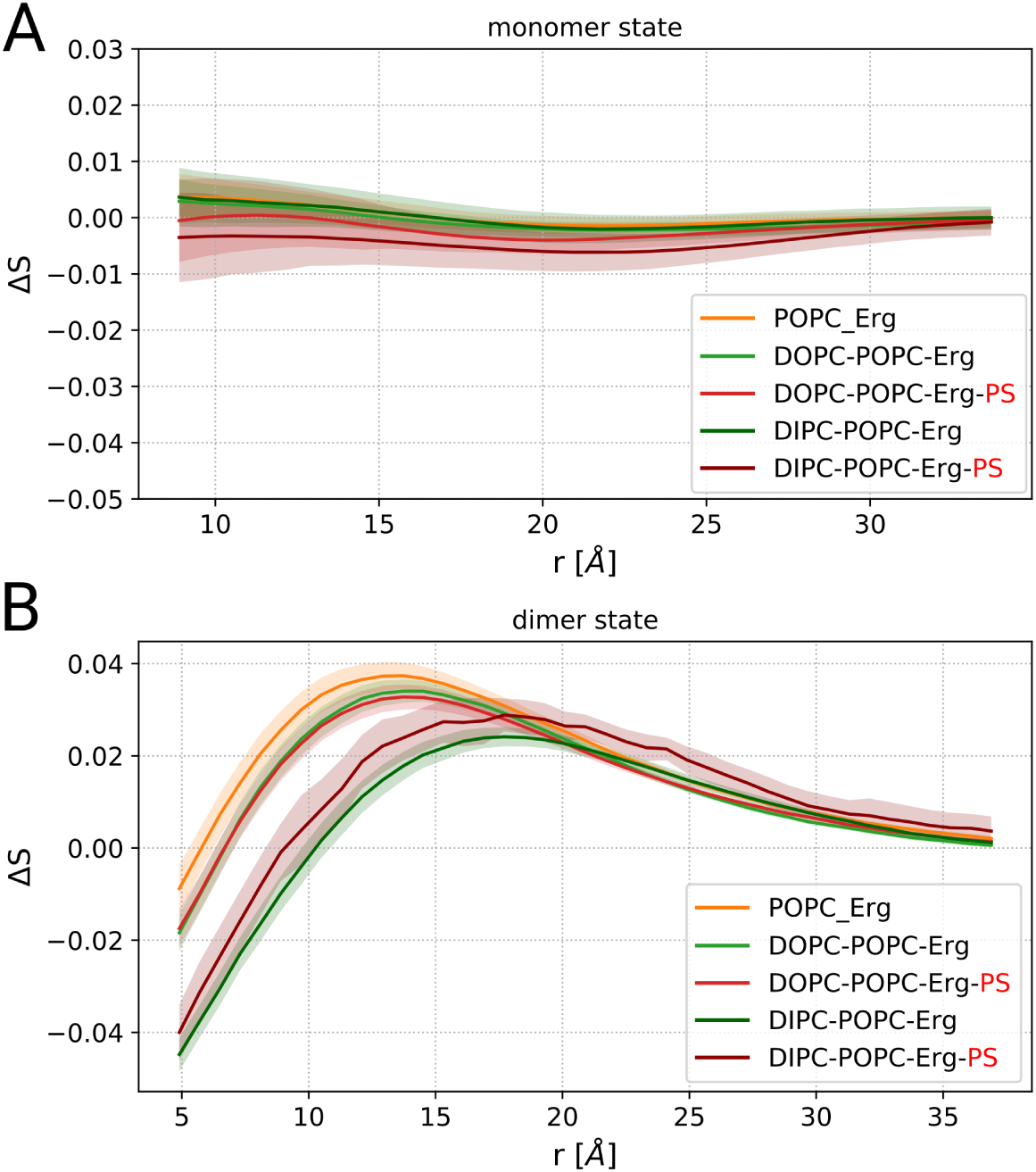
(A) The variation of order parameter of the PLs for monomer and (B) dimer state with respect to the order parameter values at large distances for the complex mixtures. In order to better visualize the effect of dimer on the membrane order, we superimposed the graphs for all systems with respect to the order parameter at large distances. The values of the order parameter at large distances for different systems are 0.39, 0.32, 0.33, 0.23, 0.24 for the respective systems listed in the legend. The shaded area represents the standard error of the mean determined using ensemble simulations.

### Experimental results

As discussed above, one way to connect atomistic simulations and mesoscopic microscopy experiments is to check the impact of a systematic variation of the lipid properties. As shown in the Methods section, we were able to experimentally control the presence of anionic PS lipids in the yeast PM and thus can check the relevance of anionic lipids.

When we tested the localization of GFP fusions to membrane segments from known single spanning yeast PM proteins, we observed correct delivery of the Slg1^TMD^ fusion protein to the PM. We then examined localization of Slg1^TMD^-mNeonGreen in more detail using a combination of TIRFM and deconvolution that we previously introduced. Slg1^TMD^ distributed in the typical network pattern (Figure 10A) that we had observed for most integral PM proteins of budding yeast. ^2^

**Figure 10:**
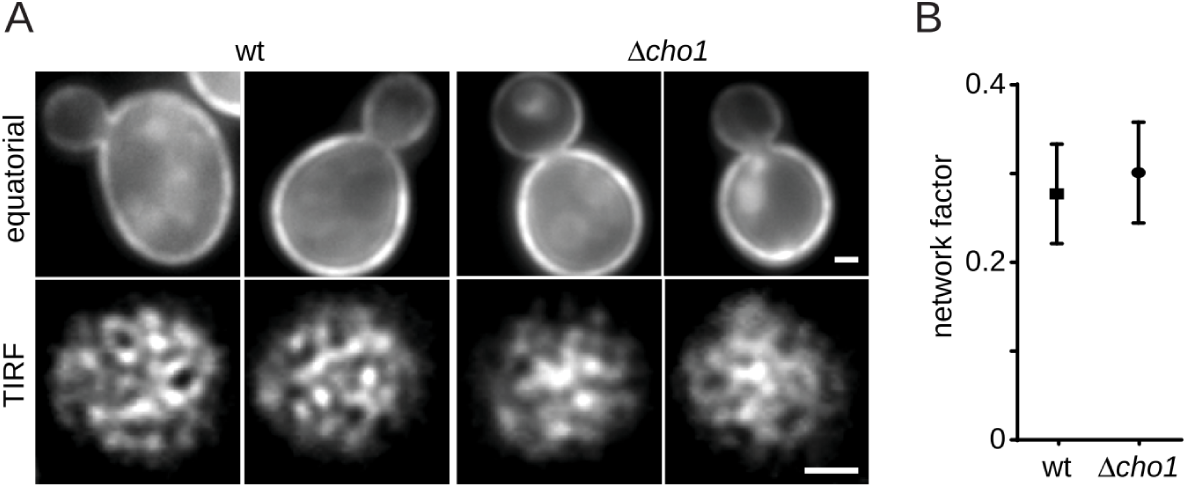
TMD distribution in the yeast PM. (A) Distribution of Slg1^TMD^-mNeonGreen in the PM seen in equatorial sections by epifluorescence (above) or at the cell surface by TIRFM (below) in control (wt) and PS-depleted (Δcho1) cells. Scale bars 1 *µ*m. (B) PM density (network factor) for Slg1^TMD^-mNeonGreen in strains shown in (A).

Our key observable, characterizing the spatial distribution of TMDs in the PM is the network factor. ^2^ In wild-type cells (in the presence of PS) the network factor of Slg1^TMD^ is around 0.27 (Figure 10B). This is in the upper range of densities observed for yeast PM proteins and represents a relatively homogeneous spatial distribution. Our results thus indicate that Slg1^TMD^ constitutes a well-suited marker to follow lateral association of TMDs in the yeast PM. Importantly, we did not observe any significant change in the network factor for Slg1^TMD^ upon depletion of PS in ÎŤcho1 cells (Figure 10A,B). This is in contrast to the large effects of PS removal that we observed for a wide range of single- and multi-spanning PM proteins (Spira 2012). However, the experimental results agree, at least on a qualitative level, with the outcome of our MD simulations. In the absence of PS lipids the binding free energy is altered by around 1 kcal/mol, with opposite signs for DOPC and DIPC. Therefore, simulations only predict a very small effect of PS on the Slg1TMD binding affinity, which is consistent with the unchanged network factor upon removal of PS.

## Discussion

We have provided detailed information about the thermodynamics and kinetics of TMD dimer formation for different lipid environments. The chosen TMD Slg1 displays a significant dimerization propensity as quantified by the free energy gain ΔG. The convergence of PMF profiles require relatively long simulation times, consistent with a recent computational study,^27^ strongly suggesting the use of a coarse-grain level of description. The chosen MARTINI force field has already provided many important insights into biological processes, in particular also about the association of TMDs.^13, 15, 19, 26, 57^ For the specific case of glycophorin A (GpA) results of the coarse-grain description ^13^ turned out to be comparable to their atomistic counterparts. ^58^ However, it has also been shown that atomistic simulations generally predict lower free energy values compared to coarse-grain ones and even underestimate the corresponding experimental data.^59^ Another study proposed that scaling values for protein-lipid interactions in atomistic simulations would result in better agreement with experimental data and better predicts the native states, while this was not the case in coarse-grain simulations. ^60^ In the present work we are more interested in relative effects upon variation of the lipid composition rather than absolute free energies.

From the simulations it was possible to distinguish the pre-dimerized and dimerized states. In the pre-dimerized state, the two TMDs are only weakly coupled and represent a LH configuration, which agrees with the observations in refs. ^13, 16, 61^ In contrast, the free energies of dimerization and the structure of the dimer depend on the lipid composition. Concerning the free energies, this dependence can be rationalized via some general characteristics. For instance, the tilt angle of the TMDs show marked differences in different systems, with DIPC giving rise to the highest tilt angle (Figure 11A). Since high tilt angles cause a reduction in the number of contacts of the N-terminus and C-terminus residues, the helix-helix association is less favorable. This correlation is also observed with respect to membrane thickness, however, all systems do not represent very distinct thickness as compared to the tilt angle of TMDs (Figure 11B). Furthermore, the free energy of association is also correlated with the membrane order parameter, reflecting the degree of saturation: the higher the order parameter, the stronger the helix-helix association (Figure 11C). The order parameter also correlates well with the tilt angle of TMDs (Figure 11A, C).

**Figure 11:**
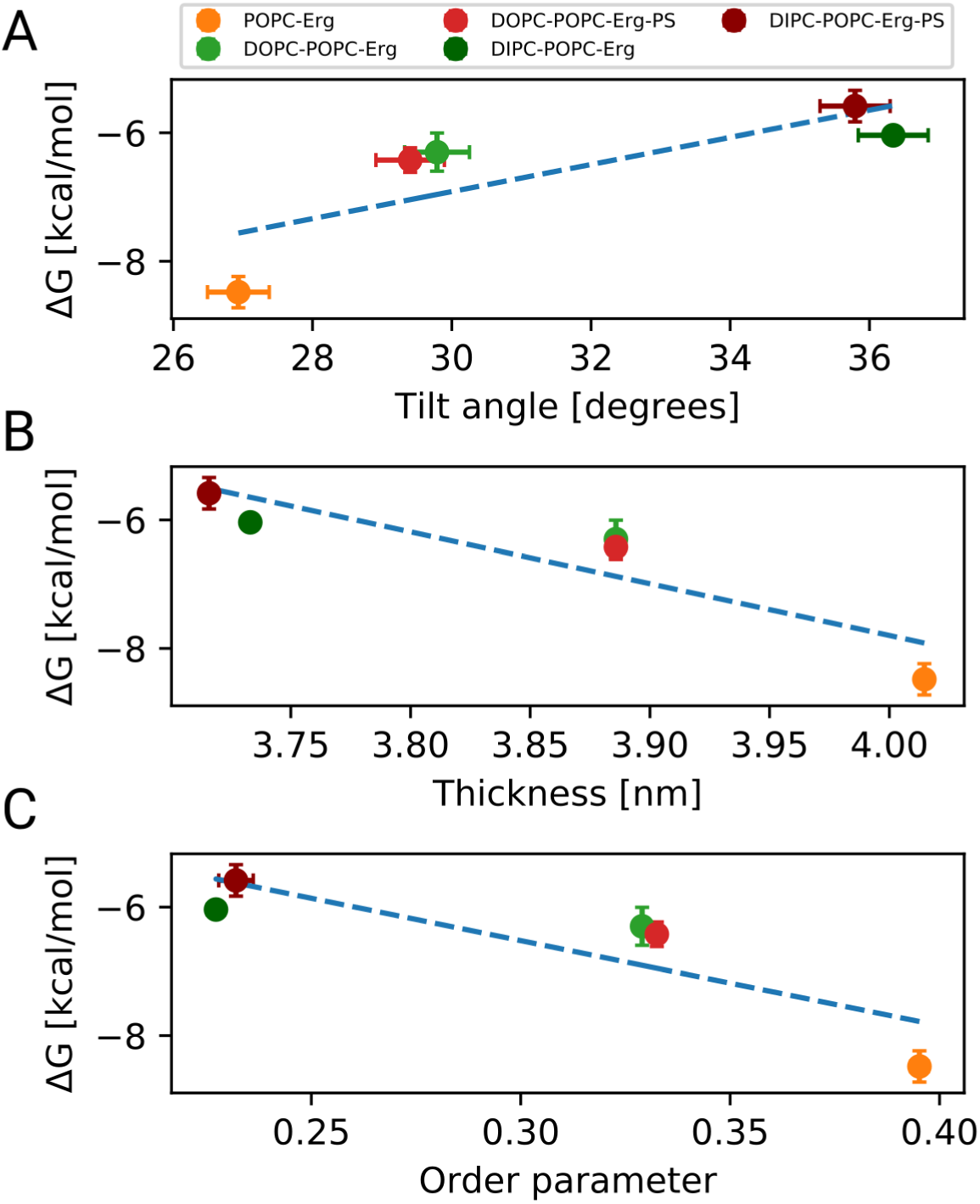
ΔG of the RH conformers in Figure 6D as a function of (A) tilt angle, (B) membrane thickness and (C) PLs order parameter.

These results partly agree with the outcome of a previous simulation study on GpA TMDs, where the correlation of dimer association with the order parameter was high. ^15^ However, no correlation with membrane thickness was observed, opposite to the results in our work. Two other aspects of that work are also worth to be compared with our results. (1). It is stated that the presence of lipids that are strongly bound to dimerized TMDs, reduce the stability of the dimer due to the loss of entropy of these lipids. Here we see that PS-lipids closely interact with the dimer due to the attraction of PS lipids to the positively-charged residues at the intracellular side of the TMDs. However, on average we do not observe a significant change of the dimer stability upon addition of PS-lipids (once slightly stabilizing, once slightly destabilizing, depending on the saturation states of the acyl chains of lipids). The GpA TMD in *E. coli* membranes, however, showed significant destabilization in the presence of anionic lipids, rationalized by the electrostatic interactions between the membrane and the positively charged residues of the protein. ^21^ Considering the similarity of the presence of the positively charged residues in the C-terminus of the GpA TMD in the aforementioned study as the studied TMD here, this difference might be due to the marked interaction of ergosterol with the TMDs in our simulations and interfering with the effect of PS lipids. (2) That work reports a small repulsion of the TMDs for intermediate distances around 2.5 nm. This is not observed in our free energy curves.

We would also like to mention a combined experimental and simulation study of bitopic proteins has shown that the free energy of dimerization is stronger in DLPC systems as compared to DPPC and POPC systems; and therefore, it was argued that neither the order parameter nor membrane thickness is correlated with dimer stability.^62^ These different observations show that a comparison between different proteins and lipid compositions have to be performed with care. Therefore, it would be interesting to incorporate different TMDs in the complex mixtures of lipids in order to understand the relevance of the proteins in lipid-protein interactions. This has been already performed in model membranes with only one lipid type and different TMDs have been shown to even exhibit opposed behavior. ^63^ The impact of cholesterol on dimer formation has already been studied for various systems. ^8–10, 27, 63^ It was also shown that ergosterol interacts with the GxxxG motif in the N-terminal part of TMDs, similar to cholesterol. ^19, 24, 26, 64, 65^ Consistently, we observed stronger interaction of ergosterol with the TMD in the upper leaflet, where these motifs reside. This strong interaction of ergosterol with TMDs may also be responsible for some of the characteristics of dimer formation, including the crossing angle state. Indeed, we saw that ergosterol interacts differently with the dimer in the POPC-Erg and DOPC systems as compared to the DIPC systems, and is a major contributor to the difference in interaction between saturated and unsaturated lipids with TMD residues (Figure 8A-D). Its higher tilt in DIPC systems increases the interaction with the residues located at the hydrophobic core of the membrane, whereas the interaction is reduced with the N- and C-terminal residues.

Of key relevance is also the presence of two different crossing angle states in dimers: RH and LH. Both states have been characterized in different experiments on ErB growth factors receptors, ^4, 66–68^ or EphA2,^69^ which are often ascribed to different dimer activity states imposed by altered membrane environment, and have also been observed in computational works. ^16, 70^ Our results also highlight the significance of membrane environment to induce different dimer states, e.g., the restriction to RH states for DIPC systems, likely responsible for various protein patterns in yeast PM. The molecular mechanism for this different behavior still needs to be revealed. The modulation of the structure of dimers by different membrane environments, comprised of only one lipid type, has also been reported for ErbB2 TMDs.^20^ Interestingly, we also showed that the free energy of dimerization for POPC-Erg and DOPC systems without PS is slightly lower for LH configurations as compared to RH ones, whereas for DOPC systems with PS the opposite holds. Of course, the resulting free energy for dimer formation has to involve an appropriate average over both the LH and the RH state.

Furthermore, the thermodynamic free-energies estimates probabilities of LH configurations were shown to be systematically smaller than the results from our ensemble simulations. Note that the ensemble simulations can only be compared to the equilibrium case after several transitions between the LH and the RH state, which would be far beyond MD time scales. The observed preference of LH configurations in the ensemble simulations thus suggest that it is more favorable to enter the LH configuration due to properties of the barrier region. This is a natural assumption since the LH configurations dominate the pre-dimerized state. To describe the complete energetic landscape of dimerization, including the structure of the dimer and saddle points of pre-dimerized states, a 2D PMF would be required, as was recently demonstrated using Metadynamics enhanced sampling technique. ^16, 27^

Interestingly, not only does the lipid environment affect TMD dimer stability, but dimerization can in turn influence the properties of the surrounding membrane. ^56^ We observed an increase in the order parameter of lipids at a distance of around 1 nm from the dimer, whereas no specific change was detected for monomers. The ordering effect was stronger for membranes with already higher order parameter. This is in agreement with stronger variation of membrane thickness around ErbB2 dimers in DPPC as compared to DLPC bilayers. ^20^ We speculate that this represents a self-reinforcing mechanism for dimer formation, since the dimerization is more preferable for systems with higher order. A similar behavior has been reported in a previous study, where dimerization of Bnip3 TMDs led to an increased order parameter in a POPC membrane. ^63^ Interestingly, for DIPC systems the lipid-protein interaction starts at a considerably larger distance than for DOPC systems, which was also reflected in the variation of the tilt angle of ergosterol with respect to the position of the dimer. Therefore, a higher degree of unsaturation in the acyl chains induces long-range perturbation in the membrane. Naturally, the change of order around a dimer may affect the mutual interaction of TMD dimers, possibly giving rise to larger clusters of TMDs.

Our results show a qualitative correlation between the mesoscopic patchwork patterns of TMDs in the yeast PM and the free energy calculations in our MD simulations. In the former approach we have examined realistic yeast PM compositions, whereas in the latter one, we have simulated simpler representative compositions, which include a number of key lipid types in actual Yeast PM. Similarly, to estimate the free energy of TMD interactions in NMR experiments, usually mimetic membranes are used. ^18, 71–73^ Interestingly, despite the wide-spread effects of PS depletion on pattern formation in the yeast PM,^2^ both MD simulations and live cell TIRF images indicated that PS lipids do not significantly influence association of the Slg1^TMD^. Future experiments will determine whether lipid saturation or other lipid varieties exert direct influence on Slg1 clustering. Furthermore, future simulations of TMDs, that show a dependence on PS levels, may help to reveal under which conditions electrostatic interactions with lipid headgroups can influence TMD dimer formation.

## Conclusions

We have studied the interaction of TMDs in a biologically relevant yeast PM, combining microscopic coarse-grain MD simulations with high-resolution microscopy experiments. Whereas the MD simulations allow a detailed characterization of local structural and thermodynamic properties, the experiments reveal aspects of the resulting structures on the 100 nm scale and beyond. We have shown that non-specific protein-lipid interactions can have significant effects on the conformation and stability of resulting dimers as well as on the free energy of dimerization. In particular, the level of acyl chain saturation was found to have an effect on TMD association. Importantly, the membrane environment is modulated upon TMD dimerization, as measured by changes in the phospholipid order parameter and the orientation of ergosterol. Dimerization in turn leads to long-range perturbations in the surrounding lipids, which penetrate further in the presence of unsaturated lipids. Our results highlight again the complex mutual interplay between the TMD and lipid environment. The role of anionic PS lipids turned out to be negligible, both observed via the free energy of association in MD simulations, and via the network factor in protein patterns in the yeast PM. Ultimately, our integrated approach aims to bridge the different scales of analysis and improve our understanding of lipid-TMD interaction during organization of biological membranes.

## Acknowledgement

We acknowledge preliminary works by Lidiya Gelemeev, atomistic simulations performed by Fabian Keller, the high performance computer resources (Palma) at University of Muenster and the financial support by the German Science Foundation (DFG) via SFB 1348 (to AH and RWS) and SFB 944 (to RWS).

## Author Contributions

AA, RWS, and AH conceived and designed the analysis. AA performed the computer simulations and the subsequent analysis. AE performed the experiments and the related analysis. All authors have contributed to the final version of manuscript.

## Supporting Information Available

The following files are available free of charge.

Figure S1. Stress profiles of the membrane. Figure S2. The autocorrelation function of the angle between two TMDs. Figure S3. Variation of the COM distance of TMDs over time; Figure S4. Average TMD-TMD contacts for the COM distance range of 10<r<20 Å; Figure S5. Overlaying average structures of dimer in coarse-grain and atomistic simulations. Figure S6. Convergence plots of the free energy profiles; Figure S7. Comparison of PMF profiles starting from bound versus unbound states. Figure S8. The distribution of crossing angle values for r=0.7 nm in bound versus unbound simulations. Figure S9. The average RDF profiles of the dimer and the lipids using a polarizable version of MARTINI force field. Figure S10. Average tilt angle of ergosterol as a function of COM distance of the dimer.

## TOC Graphic

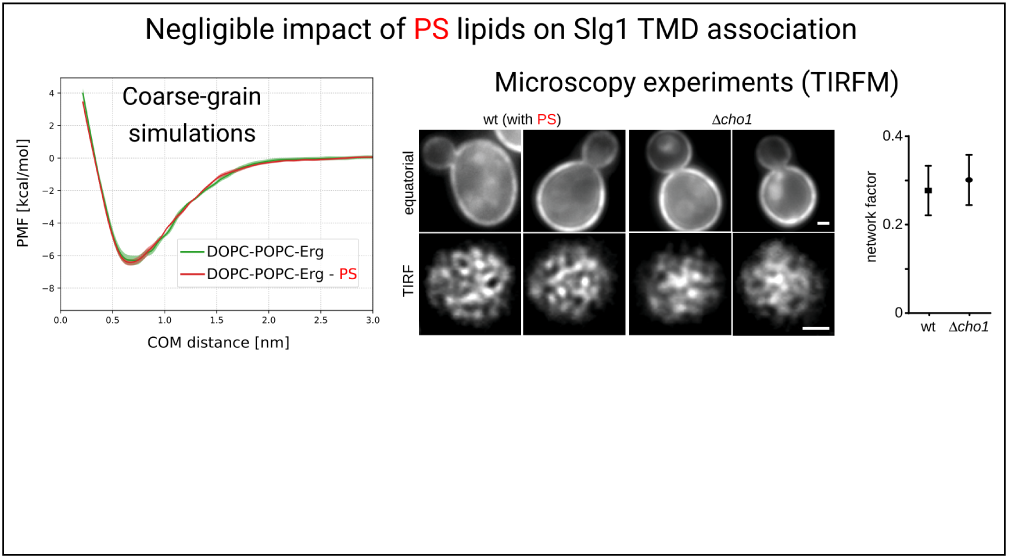

