## supplementary information for "Lipid-mediated Association of the Slg1 Transmembrane Domains in Yeast Plasma Membranes"

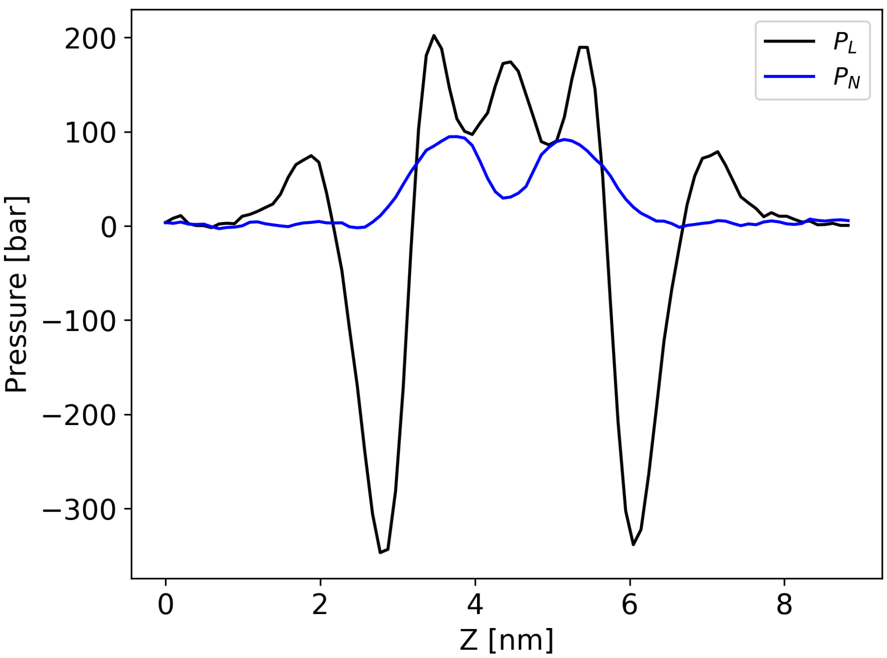


**Figure S1:** Stress profiles ($P_N = -\sigma_{zz}, P_L = -(\sigma_{xx} + \sigma_{yy})/2$) for the DOPC-POPC-Erg-PS system, as represented for one sample simulation.

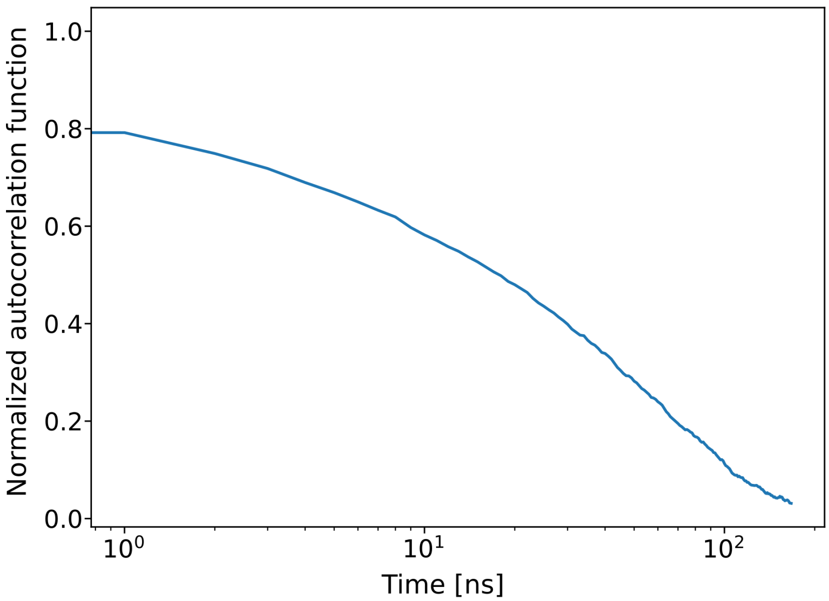


**Figure S2:** The autocorrelation function of the temporal variation of the angle between the vectors defining the orientation of the two TMDs, connecting the beginning and end of each helix for the DOPC-POPC-Erg system, averaged over all samples. The average correlation time based on the function is 32 ns.

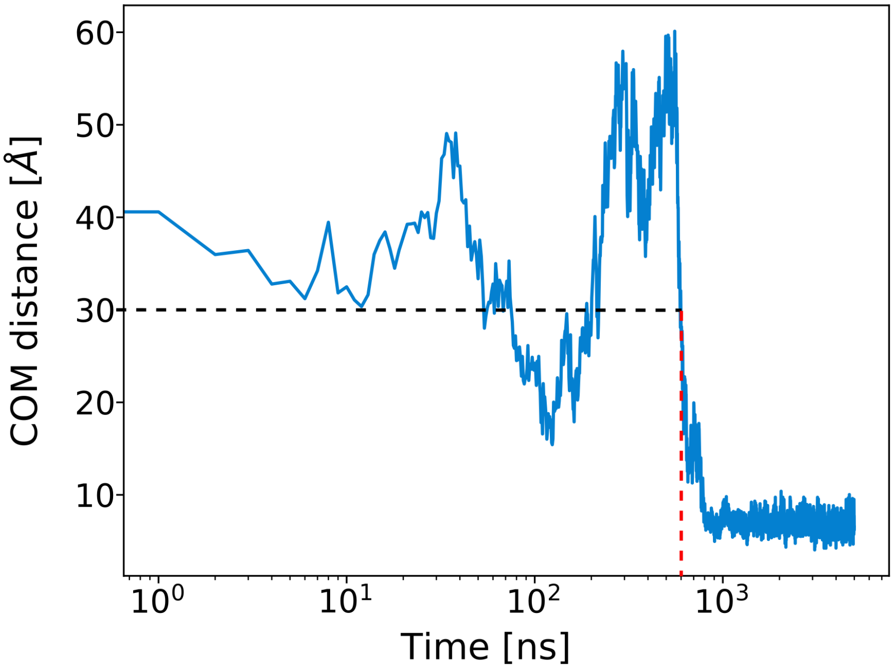


**Figure S3:** The variation of COM distance of the two TMDs during the simulation time in one sample of the DOPC-POPC-Erg-PS system is represented. The red dashed line shows the latest time at which the COM distance of the two TMDs are at the distance of 3 nm, which in this sample is around 600 ns.

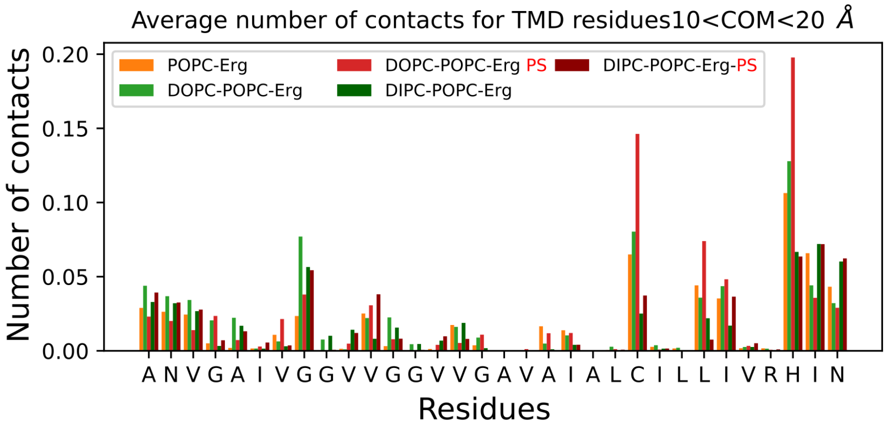


**Figure S4:** The average number of contacts between each similar pair of residues on the two TMDs is shown. A contact is considered to be formed when the distance between the COM of the two similar residues on the two TMDs is less than 6 Å.

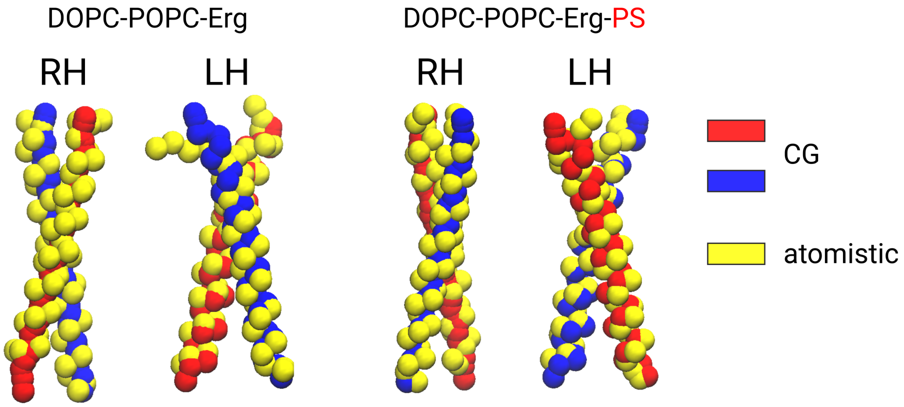


**Figure S5:** Overlay of the average structure of the RH and LH configurations of the dimer in coarse-grain simulations (red and blue) and the corresponding averaged structures in atomistic simulations, which have been back-mapped to coarse-grain, for specific COM and crossing angle values.

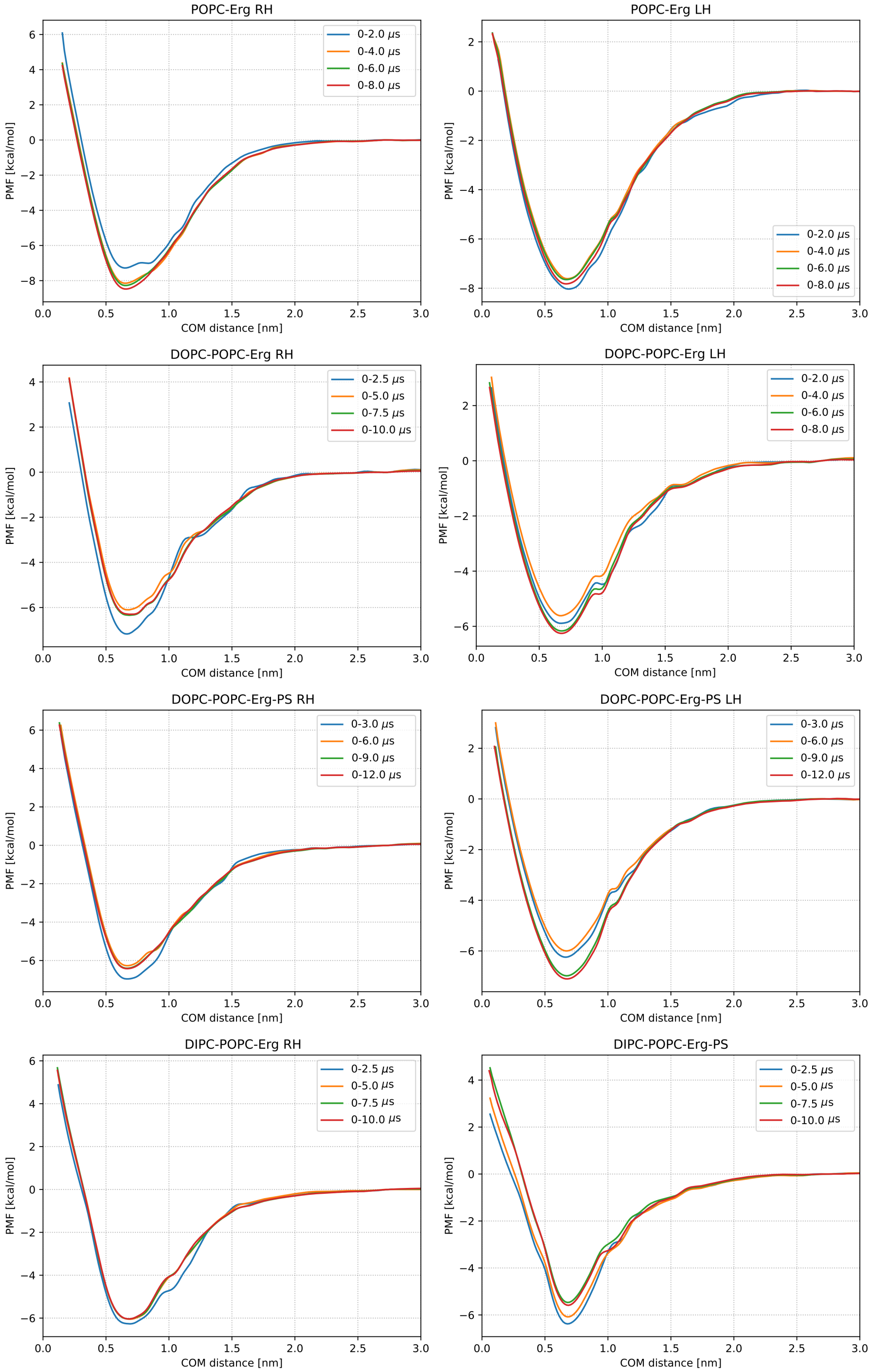


**Figure S6:** The convergence of the PMF profiles for all systems has been represented.

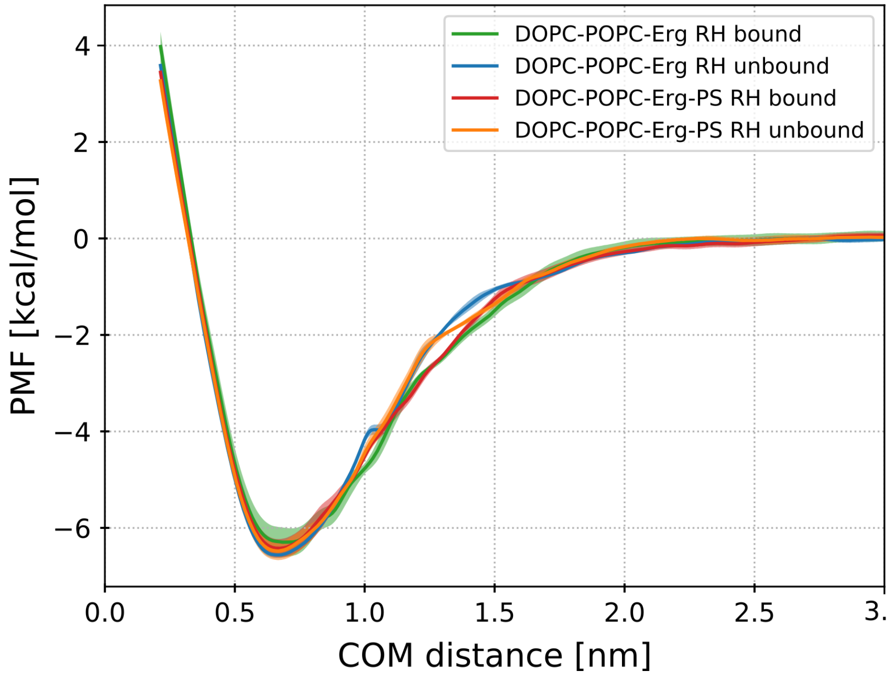


**Figure S7:** The PMF profiles for DOPC complex mixtures starting from bound and unbound initial configurations.

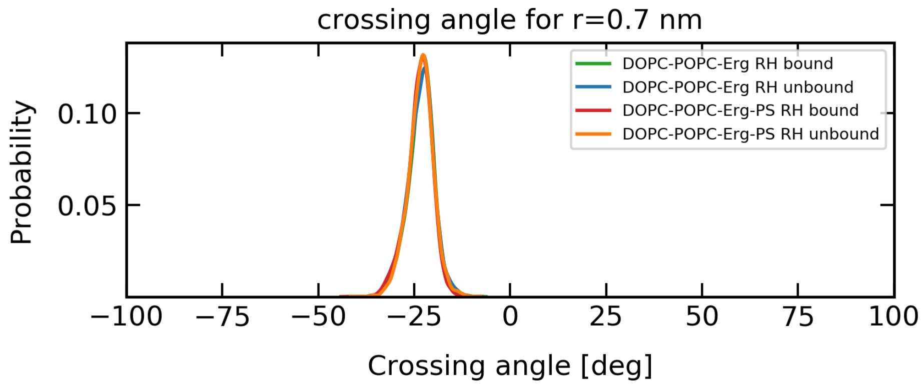


**Figure S8:** The crossing angle distribution of the dimer in the bound and unbound US simulations represented in Figure S7 for the window at which the COM distance is restrained at 0.7 nm.

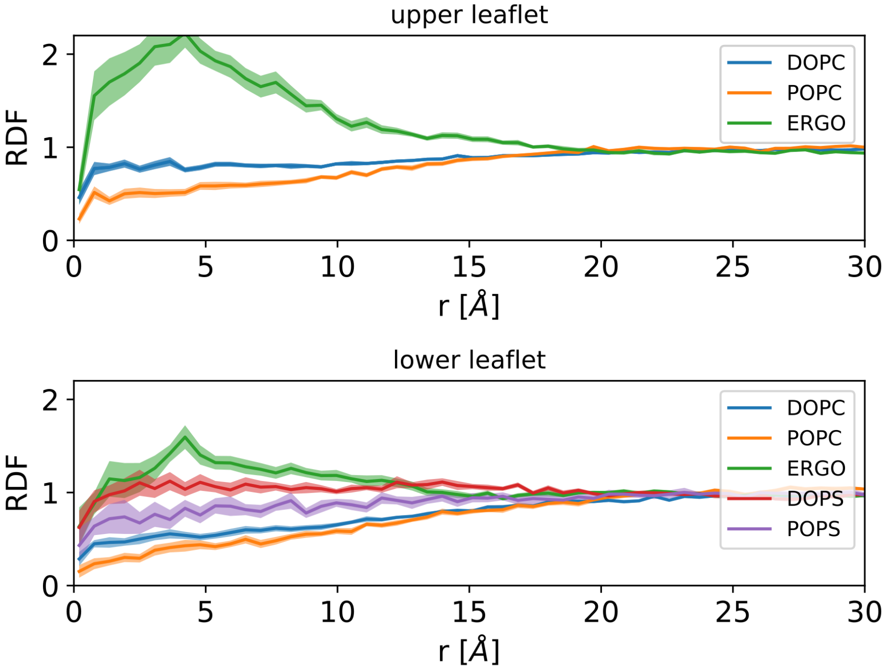


**Figure S9:** The average radial distribution function of the COM of the two TMDs and the head groups of the lipids is shown for DOPC and complex mixtures, using polarizable version of Martin force field, considering the configurations in which the dimer is formed.

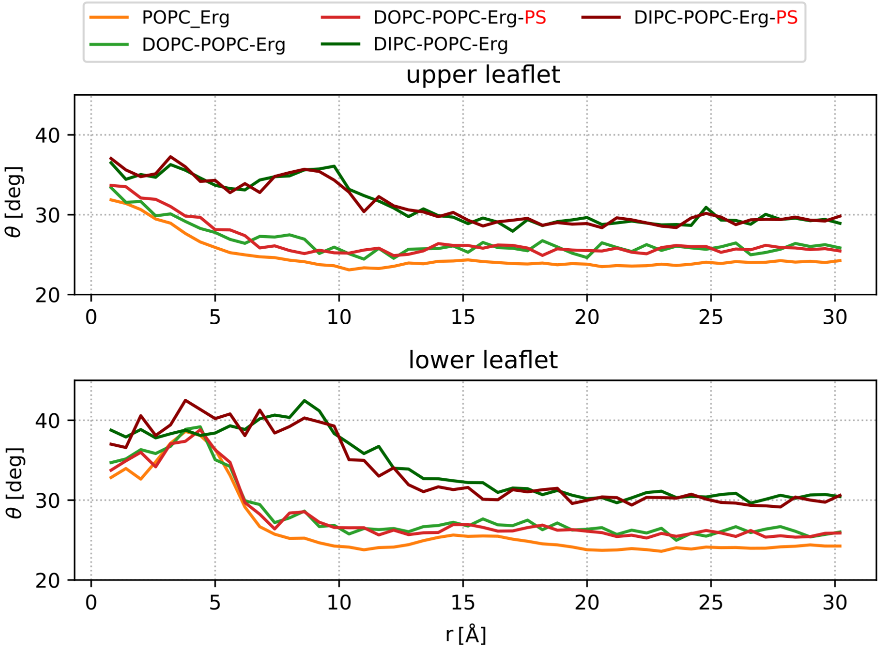


**Figure S10:** Average tilt angle of ergosterol as a function of the distance from the COM of the dimer for the upper and lower leaflet.
